# Climate change impacts on the phylogenetic diversity of the world’s terrestrial birds: more than species numbers

**DOI:** 10.1101/2020.12.02.378216

**Authors:** Alke Voskamp, Christian Hof, Matthias F. Biber, Thomas Hickler, Aidin Niamir, Stephen G. Willis, Susanne A. Fritz

## Abstract

Ongoing climate change is a major threat to biodiversity and impacts on species distributions and abundances are already evident. Heterogenous responses of species due to varying abiotic tolerances and dispersal abilities have the potential to further amplify or ameliorate these impacts through changes in species assemblages. Here we investigate the impacts of climate change on terrestrial bird distributions and, subsequently, on species richness as well as on different aspects of phylogenetic diversity of species assemblages across the globe. We go beyond previous work by disentangling the potential impacts on assemblage phylogenetic diversity of species gains vs. losses under climate change and compare the projected impacts to randomized assemblage changes.

We show that climate change might not only affect species numbers and composition of global species assemblages but could also have profound impacts on assemblage phylogenetic diversity, which, across extensive areas, differ significantly from random changes. Both the projected impacts on phylogenetic diversity and on phylogenetic structure vary greatly across the globe. Projected increases in the evolutionary history contained within species assemblages, associated with either increasing phylogenetic diversification or clustering, are most frequent at high northern latitudes. By contrast, projected declines in evolutionary history, associated with increasing phylogenetic over-dispersion or homogenisation, are projected across all continents.

The projected widespread changes in the phylogenetic structure of species assemblages show that changes in species richness do not fully reflect the potential threat from climate change to ecosystems. Our results indicate that the most severe changes to the phylogenetic diversity and structure of species assemblages are likely to be caused by species range shifts rather than range reductions and extinctions. Our findings highlight the importance of considering diverse measures in climate impact assessments and the value of integrating species-specific responses into assessments of entire community changes.

## Introduction

Global warming has been identified as one of five main anthropogenic drivers of global biodiversity loss (IPBES, 2019; Secretariat of the Convention on Biological Diversity, 2020). Whilst global warming might not represent a major threat to many species currently (Tilman et al., 2017), it is projected to increasingly threaten biodiversity in the future (Broennimann et al., 2006; Engler et al., 2011; Foden et al., 2013; Thomas et al., 2004). First responses of species to climate change have already been reported (Chen et al., 2011; Higgins et al., 2014; Jonathan Lenoir et al., 2020; Radchuk et al., 2019), covering the three possible ways in which species can adapt to global warming, i.e. through changes in their phenology, physiology or by shifting their ranges (Bellard et al., 2012). In particular, changes in species abundance and distribution have already been observed in various taxa across the globe (Bowler et al., 2017; Maclean et al., 2008; Stephens et al., 2016; Thomas, 2010). Many of these observed range shifts have been towards higher latitudes and altitudes (Hickling et al., 2006; Parmesan et al., 1999; Walther et al., 2002), but species-specific range shifts in different directions also occur (Chen et al., 2011; Moritz et al., 2008; VanDerWal et al., 2013). These idiosyncratic range shifts have the potential to be especially problematic, since they will likely result in a reshuffling of species assemblages. Potential consequences could include changes to the competitive balance between species within these assemblages (Ockendon et al., 2014) and altered predator and prey densities (Harley, 2011) as well as changes to the trait composition of local assemblages (Barbet-Massin & Jetz, 2015; Gallagher et al., 2013) and subsequently the provision of ecological functions (Pecl et al., 2017; Schleuning et al., 2020).

Compositional changes in species assemblages, caused by extinctions or range shifts, also have the potential to affect the underlying phylogenetic structure and diversity of the assemblage (Menéndez-Guerrero et al., 2020; Saladin et al., 2020). Such compositional changes can be assessed using different aspects of phylogenetic diversity. The total evolutionary diversity of a species assemblage is one such metric and measures the amount of evolutionary history that is stored within the assemblage (Flynn et al., 2011; Hardy & Senterre, 2007). Under the assumption that the evolutionary history of an assemblage indicates its evolutionary potential for adaptive change (Faith, 1992a; Forest et al., 2007), a loss of phylogenetic diversity could reduce the evolutionary potential of the assemblage, leaving it increasingly vulnerable to environmental change (Faith & Richards, 2012). This aspect of phylogenetic diversity is frequently calculated using Faith’s phylogenetic diversity (named Faith PD hereafter), which is the sum of the branch lengths of all species occurring within the assemblage back to their most recent common ancestor (Faith, 1992a). An alternative metric is the phylogenetic relatedness of the species assemblage, assessed using the mean phylogenetic distance (MPD). MPD is a measure of the deeper phylogenetic diversity of a species assemblage (Leprieur et al., 2016; Swenson & Umaña, 2014). It gives an indication of the average relatedness of the species in an assemblage. Under the assumption that closely related species have a tendency to share more similar traits than very distantly related species (Burns & Strauss, 2011), for example, an increase in the relatedness of species within an assemblage could imply a reduction in the diversity of traits present. This, in turn, could increase the vulnerability of an assemblage to environmental change (Faith, 1992a; Forest et al., 2007) but see (Jarzyna et al., 2020; Mazel et al., 2017).

Overall, MPD and Faith PD provide information on two very different aspects of the phylogenetic diversity of species assemblages; whilst the former measures the total standing evolutionary diversity across all species present (Barker, 2002), the latter measures the inverse of the average relatedness between all species pairs (Webb, 2000). Faith PD has often been found to be highly correlated to species richness for various taxa (Barker, 2002; Schipper et al., 2008), with some local exceptions where the correlation is less strong (Fritz & Rahbek, 2012; Voskamp et al., 2017). In contrast, MPD is independent of species richness (Fritz & Rahbek, 2012; Schipper et al., 2008). Since they differ mathematically, these measures have the potential to change independently of each other when species assemblages are changing (Tucker et al., 2017). Comparing these two measures when investigating temporal change in species assemblages yields additional information on the underlying phylogenetic changes that are taking place.

There are four potential directions in which the phylogenetic structure of species assemblages could shift when undergoing climate-induced compositional changes: a) MPD could increase whilst Faith PD decreases leading to increasing phylogenetic over-dispersion, i.e. increasingly distantly related species in an assemblage that represent lower values of evolutionary history; b) both MPD and Faith PD could decrease leading to increasing homogenisation; c) both MPD and Faith PD could increase leading to increasing diversification; or d) MPD could decrease whilst Faith PD increases leading to increasing phylogenetic clustering of the species assemblage, i.e. increase in clusters of closely related species that represent high values of evolutionary history.

In addition to the spatial turnover in species, changes in species richness (i.e. the gain and loss of species into and from a species assemblage) could result in non-random changes to the phylogenetic structure of species assemblages. For instance, if extinction risk is clustered across the tree of life due to similarity in species traits that confer vulnerability, this can result in the loss of entire clades and families and subsequently a disproportionate amount of evolutionary history (Russell et al., 1998). Previous studies have found this clustered pattern in Red List extinction risk assessments of taxa that include mammals, angiosperms and birds (Bromham et al., 2012; Davies & Yessoufou, 2013; Fritz & Purvis, 2010; Vamosi & Wilson, 2008). By contrast, when looking at extinction risk based on climate change projections across Europe and South Africa for the same taxa, no phylogenetic signal was found (Pio et al., 2014; Thuiller et al., 2011).

Rather than assessing the impacts of extinction across a whole phylogeny, evaluating the spatial variation in impacts of global or local extinctions on the phylogenetic structure of species assemblages could reveal how local changes in species composition impact the local phylogenetic structure. It is possible that impacts of species loss under climate change on phylogenetic diversity do not differ from a random extinction process when assessed on a global scale, but still have a strong phylogenetic impact at the local scale, weeding out entire clades from species assemblages (Huang et al., 2012). Changes in the spatial patterns of phylogenetic diversity under climate change have been shown at a regional scale under different climate change projections, for mammals, angiosperms and birds (González-Orozco et al., 2016; Pio et al., 2014; Thuiller et al., 2011). However, a study investigating potential local changes in phylogenetic diversity as caused by the projected loss of plant, mammal or insect species across the Cape of South Africa under climate change, found little difference from a random extinction process in most places (Pio et al., 2014). It is not clear to what extent such findings hold true for other taxa and more broadly across the world. Furthermore, local phylogenetic diversity under climate change will not only be subject to change through species losses but also through species that newly arrive into an area. Identifying areas where changes in species richness lead to non-random changes in phylogenetic diversity is important, since in those areas the projected changes in species richness will not reflect the entire range of impacts on the species community. For example, decreasing species richness could lead to significant homogenisation of assemblages or to significant phylogenetic over-dispersion, with very different implications for trait diversity and potential for changing competitive interactions and ecosystem function.

Here, we first (a) investigate how projected climate-induced range shifts and (local) species extinctions affect the spatial pattern of phylogenetic diversity for an entire taxon, the world’s terrestrial bird species. Secondly, (b) we compare how the projected changes in each species assemblage differ from what would be expected at random given the projected local species richness change and, for the first time, disentangle non-random changes through species entering and leaving an assemblage.

Our hypotheses are that (a), due to the high correlation between species richness (SR) and Faith PD, spatial changes in SR will be largely reflected in the spatial changes in Faith PD, whilst changes in MPD will frequently differ from this pattern. In particular, we expect Faith PD (and hence SR) and MPD to behave differently in those areas of the world where highly directional range shifts are projected to occur. Such consistent directional shifts are expected by many species at the higher northern latitudes and due to collective shifts of species towards higher altitudes (Devictor et al., 2008; J Lenoir et al., 2008; Sekercioglu et al., 2008; Virkkala & Lehikoinen, 2014). These directional shifts could potentially select for species with similar traits, leading to increasing phylogenetic clustering; this process would be identified through an increase in SR and Faith PD at the higher northern latitudes and the higher altitudes (the receiving species assemblages), accompanied by a simultaneous reduction in MPD in these areas, through related species with similar traits coming into the assemblages.

Considering the potential for non-random changes of phylogenetic diversity across species assemblages, we test a second hypothesis that (b) changes through projected species loss are decoupled from changes through projected species gain within assemblages. For example, a given assemblage could lose significant phylogenetic diversity through species loss under climate change, but also gain significantly through species gain, so the overall change in phylogenetic diversity would be marginal, though the phylogenetic structure of the assemblage could change significantly. We hypothesise that the increase in Faith PD in higher northern latitudes and higher altitudes should be lower than expected at random from the number of species gained, due to the increasing phylogenetic clustering. This clustering would also be expected to lead to significant declines in MPD (i.e. significant increase in average relatedness). Contrarily, a increase in Faith PD that is higher than expected from simply the gain of species into assemblages could occur in areas with less directional range shifts, i.e. in regions where species assemblages are more likely to be reshuffled rather than mostly gaining species. In these cases, significant changes in MPD could identify whether assemblages experience diversification (MPD increase) or phylogenetic clustering (MPD decrease). In contrast, non-random changes through the loss of species are harder to predict since they will depend on the unique evolutionary history a species brings to a local assemblage and which would be lost by its extinction or disappearance from the area. Areas that have a high number of species from ancient linages, like montane areas in tropical Africa or the northern Andes (Fjeldså & Lovette, 1997; Hughes & Eastwood, 2006), are most likely to undergo a non-random decrease in Faith PD through a loss in species richness. In such regions, significant parallel decreases in MPD would indicate strong homogenisation, whereas significant increases in MPD would indicate increasing phylogenetic over-dispersion.

## Materials and Methods

Species distribution and climatic data preparation, as well as the format for species distribution models (SDMs) and the design of the chosen dispersal buffer follow methods described in Hof et al (2018). Here, we provide an abridged summary of these methods (Hof et al., 2018), with full details in the supplementary material. The extent of our study is global, and focussed on all terrestrial areas excluding Antarctica.

### Species data

We obtained expert range maps for 9882 terrestrial bird species from BirdLife International (Birdlife International and NatureServe, 2015), considering only areas where a species was resident or occurring regularly during the breeding season. Non-breeding distributions where excluded from the analysis, because the climatic requirements of a species during the breeding season are most crucial for its survival and the non-breeding distributions of migratory species are less well known (Eyres et al., 2017; Howard et al., 2020).

### Climate data

We calculated the 19 bioclimatic variables, as described by Hijmans *et al*. (Hijmans et al., 2005), using the merged and bias-corrected meteorological forcing datasets EartH2Observe, WFDEI and ERA-Interim as provided by ISIMIP (Lange, 2016). As baseline period we used 1980 – 2009 (centred around 1995). For future projections we used the climate data provided by ISIMIP2b (Frieler et al., 2017), which comprises data from each of four different general circulation models (GCMs), i.e. MIROC5, GFDL-ESM2M, HadGEM2-ES and IPSL-CM5A-LR, for a medium warming scenario (RCP 6.0) (but see Fig. S2 to S5 and Table S2 and S3, for equivalent results based on a low warming scenario (RCP 2.6)). As a future timeframe we used end-of-century projections (2065 – 2095, centred around 2080). All climate data were provided on a 0.5° x 0.5° latitude–longitude grid.

### Species distribution models (SDMs)

We used two types of SDMs, generalized additive models (GAM, (Hastie & Tibshirani, 1990; Wood, 2006)) and boosted regression trees (GBM, (Ridgeway, 2007)), to derive the relationship between a species’ current range extent and the bioclimatic variables, following the methods in Hof et al (2018). To prepare the projected species distributions for the phylogenetic analysis, we followed the common practice of applying thresholds to transfer the projected suitability values into binary presence-absence data (Freeman & Moisen, 2008). We applied species-specific thresholds that maximized the fit to the current data, using the true skill statistic (MaxTSS) (Allouche et al., 2006). Incorporating species’ dispersal ability into future projections is vital, since the assumption of unlimited dispersal is likely to lead to unrealistic projections (Araújo et al., 2006; Berg et al., 2010; Travis et al., 2013). However, empirical natal dispersal data are available for only a very small proportion of the global terrestrial bird species (Paradis et al., 1998). Therefore, we restricted the projected future distribution of each species using estimated dispersal buffers. This approach follows previous studies that evaluated the impact of various dispersal buffers (Barbet-Massin & Jetz, 2015; Zurell et al., 2018) on projected changes in species richness, but instead of applying a constant buffer distance across all species, we applied species-specific dispersal buffers. The size of the buffer was calculated as 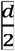, where d equals the diameter of the largest range polygon of a species (see Hof et al., 2018 for a comparison of the impact of varying sizes of dispersal buffers on species richness; and Fig. S5 to S7 and Table S4 and S5 for impacts on the projected phylogenetic measures using a more restricted dispersal assumption).

### Phylogenetic data

For the phylogenetic analysis we used the first full species-level phylogeny of extant birds (Jetz et al., 2012). We compiled a consensus tree using 150 randomly sampled trees out of the 10.000 possible tree topologies provided. For the analysis we chose the tree topologies based on the Hackett taxonomic backbone, which is the more recent of the two high-level avian topologies (Hackett et al., 2008), employed by Jetz et al. (2012). The maximum clade credibility tree topology was calculated using BEAST v1.8.4 (Drummond & Rambaut, 2007), applying the 50% majority rule and using median node heights. We matched the taxonomy used by Jetz et al. (2012) with the BirdLife version 5.0 taxonomy (Birdlife International and NatureServe, 2015), resolving all conflicting species, which resulted in a final combined dataset on the breeding range and phylogeny for 8768 species.

### Projected spatial patterns in phylogenetic diversity metrics

To extract potential changes in species richness (SR), Faith PD as well as the mean pairwise distance (MPD), we derived current and future species assemblages for each grid cell globally based on the projected species distributions.

*Change in SR* was simply calculated as the proportional change between the number of species projected to occur in a grid cell currently and in the future.

*Change in Faith PD* was calculated Faith PD for the species projected to occur in a grid cell, for both time periods, following the methods of Faith (Faith, 1992a). Faith PD is one of the most commonly used measures to calculate phylogenetic diversity (Cadotte et al., 2010). It summarizes how much of a phylogenetic tree is represented in a community by adding all branch lengths that connect the species within the community (Faith, 1992a). Change in Faith PD was then calculated as the proportional change between the current and future Faith PD value of a grid cell.

*Change in MPD* was calculated as the mean of all branch lengths that connect each pair of species within a community (Clarke & Warwick, 1998; Kembel et al., 2010; Webb et al., 2002). It gives an indication of how closely related individuals are, on average, within a community (Tucker et al., 2017). Subsequently, change in MPD was again calculated as the proportional change between the current and future MPD value of a grid cell.

### Projected non-random changes in phylogenetic assemblage structure

We evaluated whether the projected changes in Faith PD and mean pairwise distance (MPD) were different from what could be expected if the species that moved in or out of an area were randomly distributed across the phylogeny. These randomizations are necessary because the structure of the phylogeny determines the possible extent of projected changes in Faith PD and MPD given a particular species assemblage and number of species moving in or out (May, 1990; Purvis et al., 2000). We decomposed the net change in SR in a given assemblage (grid cell) into the species persisting under climate change, the species projected to be lost (through extinction or emigration), and the species projected to be gained (through colonisation) under climate change (Fig. 1).

**Fig. 1:**
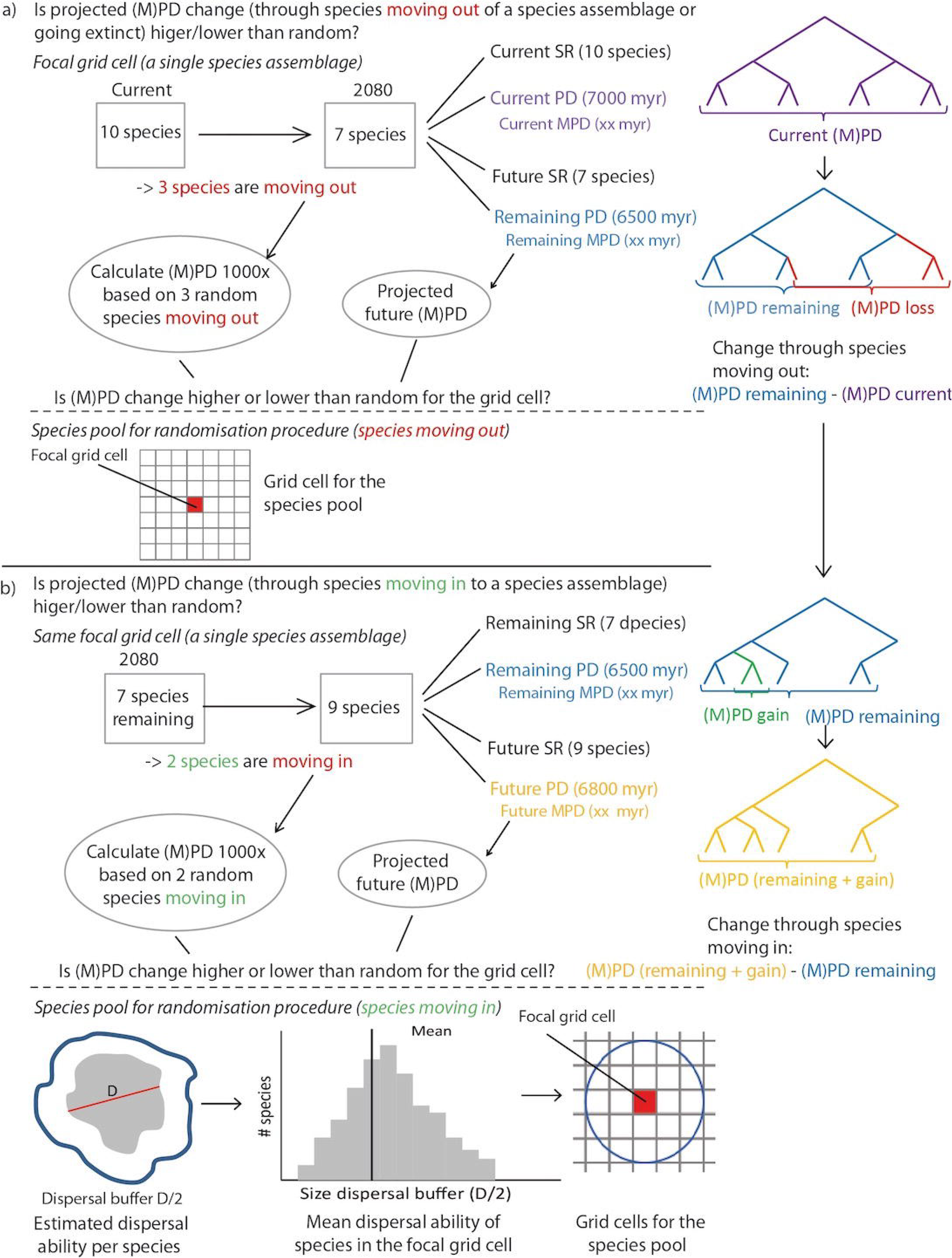
Methods used to compare the projected changes in Faith PD (called PD in the flow diagram) and mean phylogenetic distance (MPD) of a species assemblage (grid cell), based on species that are projected to be lost from (a) and gained into (b) the assemblage, with the expected changes in Faith PD/MPD based on the same number of species being lost and gained at random. In this example, we assume that a) there were 10 species in the assemblage initially and we project 7 species to remain in the assemblage (with 3 species projected to emigrate or go extinct). To calculate the expectation for random species loss, we then drop 3 random species from the list of ten species 1000 times, i.e. the species pool is just the focal assemblage in this case. We then recalculate the two metrics of phylogenetic assemblage structure for each random assemblage, and compare this expected change under random species loss to the projected change in the two metrics. Then, we assume that b) seven species remain in the assemblage and two species are projected to be gained to the assemblage. To calculate the expectation for random species gain, we draw 2 random species from the species pool 1000 times, where the colonist species are drawn randomly from a pool of candidate species that occur within a colonisable distance of the focal assemblage; the mean dispersal ability for each species is estimated as half the value of the longest range diameter (D). We then recalculate the two metrics of phylogenetic assemblage structure for each random assemblage, and compare this expected change under random species gain to the projected change in the two metrics.

Random changes in both phylogenetic measures of a species assemblage (grid cell), through species loss, were calculated using the list of species projected to be currently present in the assemblage. The same number of species as projected to be lost (by the SDMs) from the assemblage was then repeatedly (1000 times) dropped from the current species assemblage at random, and both phylogenetic measures were recalculated each time (Fig. 1a). The change in Faith PD or MPD was then calculated as Faith PD_remaining_ *minus* Faith PD_current_ or MPD_remaining_ *minus* MPD_current_, respectively (Fig. 1a). Finally, based on the 1000 repeats we calculated a two-sided p-value as the proportion of random values that were smaller or larger than the observed value. This p-value indicates if there was a significant difference between the projected change in the phylogenetic measures and the changes based on random species removals.

When projecting potential changes to species assemblages under climate change, there will not only be species that are lost from the assemblage, but also species that are gained by the assemblage (Fig. 1b). To calculate if the change in both phylogenetic measures, based on species projected to be gained by the assemblage (colonising the assemblage), was different from what one would expect if species would have been gained at random, we again compared the lists of species IDs projected to occur in the assemblage currently and in future. We extracted the number of species that are projected to be gained by the assemblage (grid cell) and then, using this number, we randomly added new species to those that were projected to remain in the assemblage under climate change, using a species pool defined based on estimated species’ dispersal abilities as explained in more detail below (Fig. 1b). The change in Faith PD or MPD was then calculated as Faith PD_(remaining + gain)_ *minus* Faith PD_remaining_ or MPD_(remaining + gain)_ *minus* MPD_remaining_, respectively. Again, we calculated a two sided p-value indicating if there was a significant difference between the projected change and the random change in both phylogenetic measures based on random species being gained by the assemblage.

Drawing random species to be gained by an assemblage (grid cell) from a species pool containing the whole list of terrestrial birds included in the analysis (8269 globally distributed species) would yield highly unlikely results, because many species would be unable to move into the area due to climate or habitat requirements, dispersal ability or dispersal barriers. To produce more realistic projections, we created assemblage-specific (grid cell-specific) species pools to draw the species from (Fig. 1b). To create these species pools we used the estimated species-specific dispersal buffers we applied for the projections (see SDM methods). For each species assemblage we extracted the mean estimated dispersal distance across all species occurring within the assemblage, to have an estimate on how far birds are projected to disperse within this area. We then used this mean distance to create a buffer around the species assemblage (grid cell). The species pool for an individual assemblage subsequently contained all the species that occurred within this buffer. We used these assemblage-specific dispersal buffers because the average natal dispersal distance varies globally, with on average much shorter dispersal distances in the tropics (Janzen, 1967; Salisbury et al., 2012).

The final dataset containing the species assemblage values needed to run the analysis and create the plots can be found on Zenodo (10.5281/zenodo.4262462). The R code for the analysis is provided on GitHub (https://github.com/AlkeVoskamp/Climate_change_PD_MPD.git). *BOTH WILL BE MADE PUBLIC UPON ACCEPTANCE*

## Results

### Projected spatial patterns in phylogenetic diversity metrics

The projected changes in species richness (SR) within assemblages (grid cells) do not differ greatly across continents but do differ within them (Fig. 2a). These SR changes are spatially highly correlated with projected changes in Faith PD across the globe (Fig. 2b). Although proportional losses in both SR and Faith PD are likely to be most extreme in species-poor regions (e.g. deserts of Middle East, Sahara, Australia and southern Africa), we also project high proportional losses in some of the very species-rich regions of the world (Fig. 2d and 2e). Across the species-rich regions the proportional changes in Faith PD and SR are especially severe in parts of South America, such as the Amazon region, Uruguay and northern Argentina, as well as on New Guinea, but assemblages with losses up to 30 % can be found across all continents (Fig 2d and 2e). Assemblages with a projected proportional gain in both measures are especially widespread at high northern latitudes across the Nearctic and Palearctic realm (Fig 2d and 2e).

**Fig. 2:**
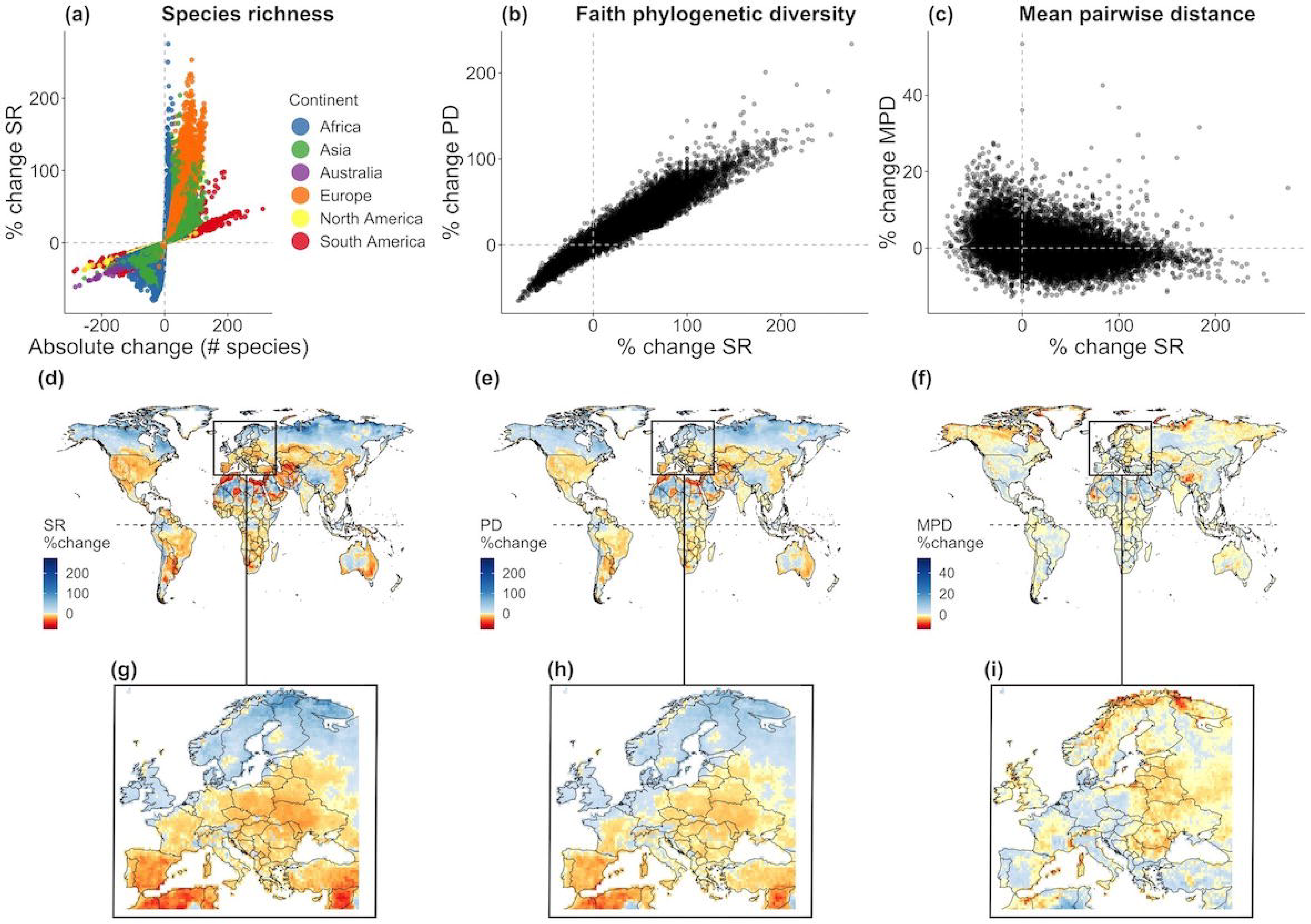
Projected changes in species richness (SR), Faith’s phylogenetic diversity (Faith PD) and mean phylogenetic distance (MPD) under a medium emission scenario (RCP6.0) and a medium dispersal scenario by 2080. (a) shows the percentage change in SR against absolute change in SR; (b) the percentage change in Faith PD against percentage change in SR; (c) the percentage change in MPD against percentage change in SR (d) the spatial distribution of percentage change in SR; (e) the spatial distribution of percentage change in Faith PD and (f) the spatial distribution of percentage change in MPD. The percentage change for all three measures is shown in detail for Europe (g – i). Red indicates a negative change (e.g. loss in species richness, Faith PD or MPD), blue indicates a positive change (e.g. gain in species richness, Faith PD or MPD).

The projected changes in mean phylogenetic distance (MPD) differ substantially from the projected changes in SR (Fig. 2c) and Faith PD. Looking at the same areas described above for changes in SR and PD, we find, that assemblages for which we project a decrease in MPD are widespread across the northern Nearctic and Palearctic (Fig. 2f), whereas assemblages projected to experience an increase in MPD are located in the southern parts of the Amazon, Uruguay and northern Argentina as well as New Guinea. Overall, the spatial patterns of projected change in MPD are often opposite to those in Faith PD and SR (Fig. 2d-f). An example for this opposite trend in the three indicators are the changes projected across Europe. Increases in Faith PD (and gains in SR) are mainly projected in the northern parts, across the UK and Scandinavia, whereas decreases are projected to be widespread across mainland Europe including Spain (Fig. 2g-h). On the contrary, Scandinavia and the UK are projected to experience decreases in MPD, whereas increases are scattered across mainland Europe including Spain (Fig. 2i).

Overall, when looking at projected changes in the two phylogenetic structure metrics (Fig. 2), we find that Faith PD and MPD are changing into opposite directions in approximately 60% of the global terrestrial area (Table 1). Exploring the correlation and divergence between Faith PD and MPD further (Fig. 3, Table 1), we find that 30% of the global species assemblages (grid cells) are projected to experience an increase in MPD and a decrease in Faith PD, i.e. a loss in average relatedness and a loss in standing evolutionary history, leading to increasing phylogenetic over-dispersion of these species assemblages. In 15% of the species assemblages globally, both MPD and Faith PD decrease, leading to increasing relatedness and decreasing evolutionary history, i.e. an increasing homogenisation of these species assemblages. Further, 29% of the species assemblages experience a decrease in MPD and an increase in Faith PD, indicating increasing relatedness and evolutionary history which leads to phylogenetic clustering of the species assemblages. Finally, 26% of the species assemblages are experiencing increases of both MPD and Faith PD, indicating losses of relatedness and gains in evolutionary history that translate to overall increasing diversification of these species assemblages under global warming.

**Table 1:** The overall terrestrial area, globally and per continent, that falls into the four different categories of combined change in two phylogenetic structure metrics, Faith’s phylogenetic diversity (Faith PD) and mean phylogenetic distance (MPD) (as shown in Fig. 3): increasing homogenisation (loss of PD and MPD); Increasing clustering (gain in PD and loss of MPD); Increasing over-dispersion (loss of PD and gain in MPD) and Increasing diversification (gain in PD and MPD). The area extent is given in km^2^ as well as in the percentage of the total terrestrial area, per continent and globally. The extent of the area projected to fall into the four different categories is derived assuming a medium emission scenario (RCP6.0) and a medium dispersal scenario by 2080.

|  | **Increasing homogenisation** |  | **Increasing clustering** |  | **Increasing over-dispersion** |  | **Increasing diversification** |  |
| --- | --- | --- | --- | --- | --- | --- | --- | --- |
|  | *Area in km<sup>2</sup>* | *%* | *Area in km<sup>2</sup>* | *%* | *Area in km<sup>2</sup>* | *%* | *Area in km<sup>2</sup>* | *%* |
| **Africa** | 4,321,971 | 15 | 5,572,879 | 19 | 13,159,337 | 44 | 6,813,364 | 23 |
| **Asia** | 4,928,362 | 17 | 6,055,430 | 29 | 12,751,917 | 42 | 7,307,249 | 23 |
| **Australia** | 2,345,390 | 28 | 1,830,899 | 22 | 2,834,075 | 33 | 1,467,003 | 18 |
| **Europe** | 3,223,093 | 12 | 7,390,459 | 37 | 2,937,947 | 10 | 9,160,714 | 41 |
| **North America** | 2,969,140 | 12 | 7,371,192 | 41 | 7,060,167 | 23 | 5,383,293 | 25 |
| **South America** | 1,603,745 | 9 | 3,829,585 | 22 | 9,151,380 | 52 | 3,015,050 | 17 |
| **Global km<sup>2</sup>** | 19,391,701 |  | 32,050,444 |  | 47,894,823 |  | 33,146,674 |  |
| **Global %** | 14 |  | 29 |  | 31 |  | 27 |  |

**Fig. 3:**
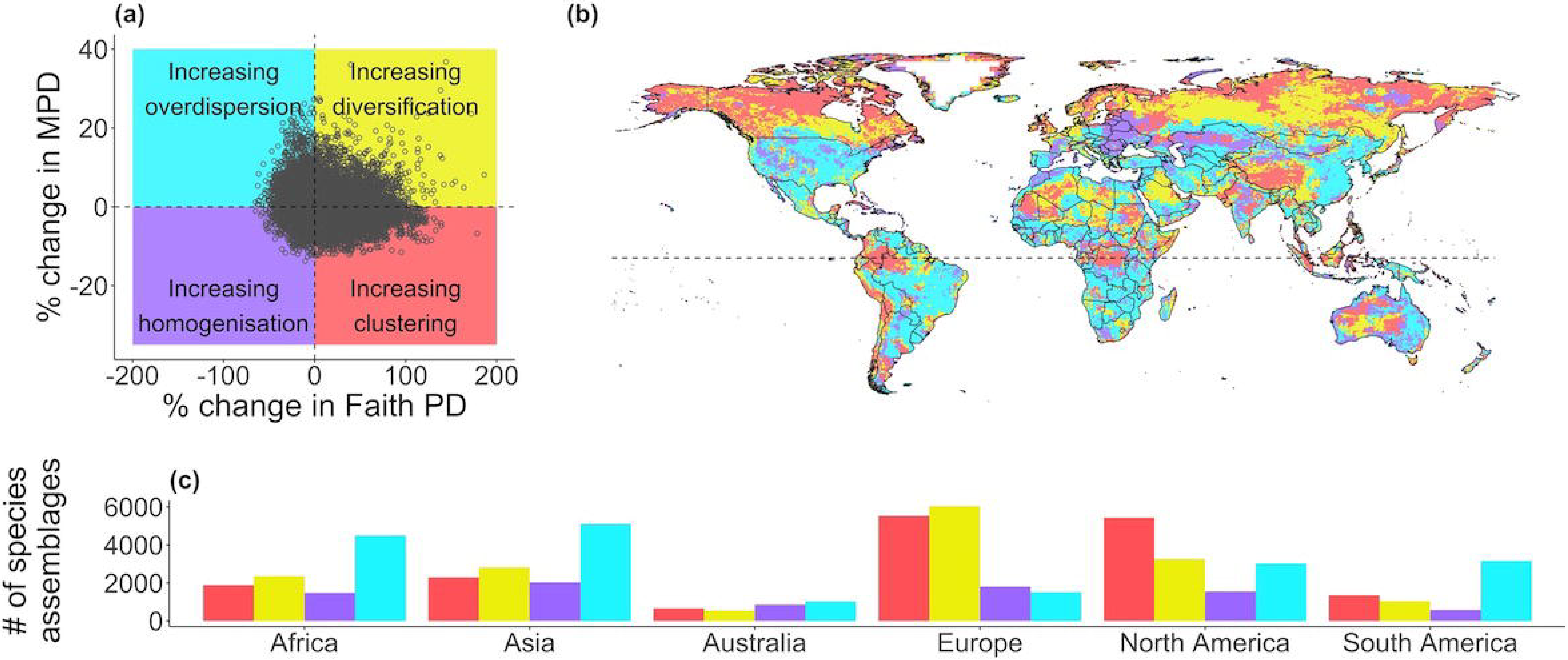
Comparison of the direction of projected changes in phylogenetic assemblage structure as indicated by mean phylogenetic distance (MPD) versus by Faith’s phylogenetic diversity (Faith PD) under a medium emission scenario (RCP 6.0) assuming a medium dispersal scenario by 2080. The scatterplot (a) shows percentage change in MPD against percentage change in Faith PD, divided into four categories of change using the median along each axis. The map (b) shows the spatial distribution of the species assemblages falling into one of these four categories, and the bar chart (c) shows the number of assemblages per category across different continents. The four defined categories are: grid cells with a projected gain in MPD and loss in Faith PD leading to increasing phylogenetic overdispersion of these species assemblages (blue); grid cells with a projected loss in both MPD and Faith PD, leading to increasing homogenisation of these species assemblages (purple); grid cells with a projected loss of MPD and gain in Faith PD, indicating increasing phylogenetic clustering of these species assemblages (red); and grid cells with a projected gain in both MPD and Faith PD, indicating increasing diversification within these species assemblages (yellow).

### Projected non-random changes in phylogenetic assemblage structure

We identify areas where the projected changes in each phylogenetic diversity metric, i.e. Faith PD and MPD, are higher or lower than we would expect from the projected changes in SR, by randomising the identity of species that were gained by or lost from a species assemblage. Focussing only on the areas where the projected changes differ significantly from what would be expected at random, based on the two-sided p-value, we find that areas where the decrease in Faith PD (through the loss in SR) was significantly less than expected from randomized species moving out of the assemblage occur on all continents but are most frequent in the northern Palearctic and Nearctic (Fig. 4a). In these areas, assemblages are projected to lose species through climate change that represent unusually low amounts of evolutionary history. In contrast, areas with a significantly stronger decrease in Faith PD than we would expect at random (through the loss of species) also occur on all continents but are most common in central South America and southern African regions (Fig. 4a). These areas are therefore projected to lose species that represent disproportionately high amounts of evolutionary history in their respective assemblages.

**Fig. 4:**
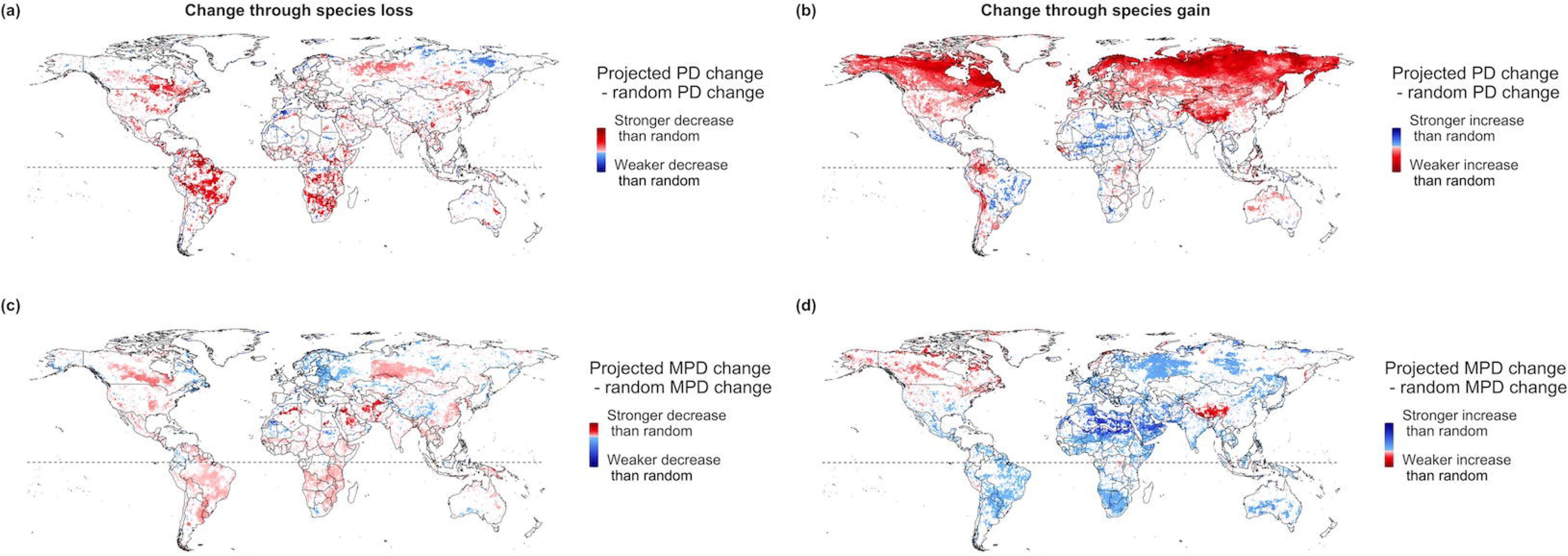
The significance and direction of projected changes in Faith’s phylogenetic diversity (Faith PD) and mean phylogenetic distance (MPD) of species assemblages (grid cells), through species that are projected to be lost from (a and c) and gained into (b and d) assemblages, in comparison to expected changes if species were lost and gained at random. Difference values for species being lost from an assemblage are calculated as shown in Fig 1. For the maps of change in Faith PD/MPD through species being lost from an assemblage (a and c), red indicates that the loss of Faith PD/MPD caused by the species that are projected to be lost from the assemblage is significantly higher than what would be expected if the same number of random species would be lost; blue indicates that the loss is significantly lower than what would be expected if random species would be lost (significance is derived using a two-sided p-value < 0.05 or > 0.95). For the maps of change in Faith PD/ MPD through species being gained into an assemblage (b and d), red indicates that the gain in Faith PD/MPD through the species projected to be gained into the assemblage is significantly lower than what would be expected if the same number of random species would be gained into the assemblage, blue indicates that the gain is significantly higher than what would be expected if random species would be gained. A gain or loss in Faith PD signifies a significant increase or decrease in total evolutionary history represented, respectively; a gain or loss in MPD signifies a significant decrease or increase in average relatedness, respectively. White areas in each map have no significant changes compared to random species gain or loss. Results are shown for a medium emission scenario (RCP6.0) and a medium dispersal scenario by 2080.

Focussing on the increases, we find that areas with significantly lower increases in Faith PD than would be expected through equivalent random species gains are most frequent at high northern latitudes, stretching across the entire Nearctic and Palearctic realm, but also across parts of South America and Australia (Fig. 4b). This category of significantly lower increase in Faith PD is the most widespread in extent, indicating that not only are more areas projected to gain more species than lose them (cf. Fig. 2d), but also that the species gains in these areas do not lead to the expected increases in evolutionary history in a large part of the world. Areas with a significantly higher increase in Faith PD than expected by chance (through the gain in SR) are much less common and are mainly located in northern Africa, northern central America and the eastern half of South America (Fig. 4b). These areas are projected to gain species that represent unusually high amounts of evolutionary history, potentially making those assemblages more diverse and increasing their evolutionary potential.

For the changes in MPD, we find areas where the projected loss in SR leads to significantly lower declines in MPD than equivalent random losses occur globally, but are generally most extensive in the Nearctic and Palearctic realm (Fig. 4c). In these areas, mean relatedness either decreases or does not increase as much as expected from the projected species loss, indicating that assemblages are projected to lose species that were significantly less closely related to remaining species than expected under random loss. In contrast, the projected loss in SR leads to a significantly stronger decrease in MPD than expected at random in Northern Africa, Arabia and the Middle East, but also to significant but less strong decreases across the Americas as well as in southern Africa and west Siberia (Fig. 4c). In these areas average relatedness of species within assemblages either increases significantly more, or declines significantly less, than expected under random species loss; this suggests that the species projected to be lost are disproportionately more closely related to the remaining species.

Similar to the projected changes in Faith PD, the non-random changes in MPD are much stronger from species gain under climate change (Fig. 4d) than from species loss (Fig. 4c). Areas where the increase in SR leads to significantly less increase in MPD than expected from equivalent random species additions are nearly completely restricted to the Nearctic realm as well as the northern Himalayas and Qinghai-Tibet Plateau (Fig. 4d). In these areas, assemblages tend to gain species that are significantly more closely related to existing species than expected at random. Areas where the increase in MPD is significantly higher than would be expected by an equivalent gain in random species are widespread across most continents (Fig. 4d), indicating that average relatedness across most of the world decreases through species projected to be gained under climate change, i.e. that assemblages gain species that are more distantly related to existing species than expected at random.

When comparing the significant non-random increases vs. decreases in phylogenetic diversity within each assemblage, we find that there are numerous assemblages where projected significant changes in phylogenetic structure overlap, e.g. stronger decreases than expected from random species loss and, at the same time, weaker increases than expected from random species gain (Table 2). Projected future changes in Faith PD that differ significantly from equivalent random species losses or gains overlap most in assemblages that show a stronger decrease and coinciding stronger increase of evolutionary history than expected, i.e. these assemblages will lose more phylogenetic diversity than if species were lost randomly and simultaneously gain more phylogenetic diversity than if species were gained randomly (>14% of global assemblages; Table 2, second row). This indicates that particularly high proportions of assemblages in most continents (in South America, Africa, North America and Europe) are projected to experience species reshuffling through both species losses and gains. Our results are similar for MPD, where >14% of global assemblages are projected to decrease significantly more than under random species loss and simultaneously increase significantly less than under random species loss, indicating that the species losses and gains will both cause an unusually strong increase in net relatedness of species in these assemblages (Table 2, second row; particularly in South America, Africa, North America, and Asia). For Faith PD, the only other category of coinciding significantly non-random changes is that of simultaneous stronger decrease and weaker increase (Table 2, first row), indicating that >10% of assemblages in Europe and North America are projected to experience unusually strong decreases in Faith PD and therefore lose significant amounts of evolutionary history through both species losses and gains. For MPD, a vast majority of assemblages does not experience any of the overlap categories, with the exceptions of the above-mentioned ones, although nearly 10% of European assemblages are projected to experience simultaneous stronger increases and stronger decreases in relatedness than expected under random species loss and gain (Table 2, third row). This further corroborates the results for Faith PD, indicating that a high proportion of assemblages in Europe stand to experience major species reshuffling.

**Table 2:**
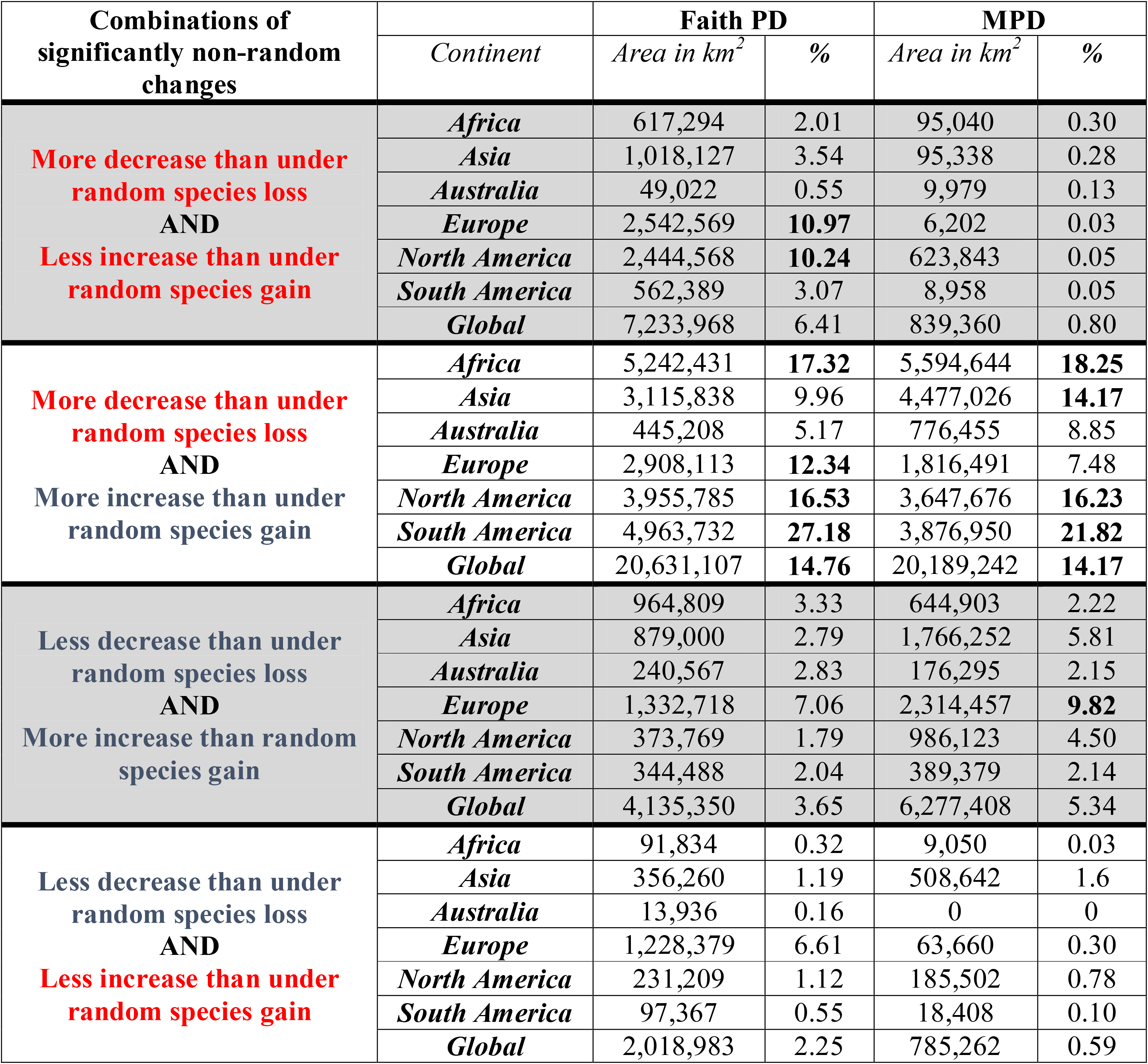
Combined projected changes in Faith’s phylogenetic diversity (Faith PD) and mean phylogenetic distance (MPD), indicating the proportions of those assemblages across the globe where each measure changes significantly compared to both, a randomized gain and a randomized loss of species (as shown in Fig. 4). The area extent is given in km^2^ as well as in terms of percentage of the total global land mass; percentage values above 10% are printed in bold. The extent of the area projected to fall into the four different combinations is derived assuming a medium emission scenario (RCP6.0) and a medium dispersal scenario by 2080.

## Discussion

We found that the projected impacts of climate change not only affected the amount of evolutionary history stored within species assemblages but also had significant impact on the phylogenetic structure of species communities. The independent changes in the two phylogenetic diversity measures, Faith PD and MPD, resulted in four main directions in which the phylogenetic structures of species communities can change. The frequency with which these structural changes occurred showed a latitudinal gradient, with increases in the amount of evolutionary history contained within species assemblages, associated with either increasing phylogenetic diversification or clustering, being most frequent at high northern latitudes. By contrast, declining evolutionary history, associated with increasing phylogenetic over-dispersion or homogenisation, occurred across all continents. Overall, the projected changes in Faith PD and MPD differed significantly from what we would expect if random species were being gained or lost across large areas of the globe, indicating that the phylogenetic assemblage structure might be changed completely in the future through local species loss and gain that is strongly selective in terms of phylogeny.

### Projected spatial patterns in phylogenetic diversity metrics

As expected, our results showed that the spatial patterns of proportional changes in species richness (SR) and Faith PD are highly correlated on a global scale (Fig. 2b), thus the projected losses as well as gains in assemblage SR is largely reflected in their decreases and increases in Faith PD (Figure 1). By contrast, the changes in mean phylogenetic distance (MPD) are independent from the changes in SR and Faith PD and frequently show an opposite pattern. These striking differences are corroborated by the strong spatial patterns in those changes that are significantly non-random (Fig 4, Table 2, supplementary discussion).

Focussing on the example of Europe (Figure 2), the increase in Faith PD in combination with the simultaneous decrease in MPD across the UK and Scandinavia indicates that, although these areas tend to gain species, the projected future assemblages might comprise more closely related species in the future than currently. The projected gains in species in these areas are in line with already observed northwards shifts of terrestrial bird species (Thomas & Lennon, 1999; Virkkala et al., 2014). The projected simultaneous decrease in MPD in these areas supports the idea that species responses to climate change might in some cases be linked to species traits, with species that have similar traits showing similar responses (Leach et al., 2015; Mason et al., 2019). If these traits are clustered across the phylogeny, which is the case for some traits (Barnagaud et al., 2014; Böhning-Gaese & Oberrath, 1999); but see (Khaliq et al., 2015; Losos, 2008)), this could explain the projected gain in species richness and increase in Faith PD as well as the simultaneous increase in relatedness within northern species assemblages.

A decrease in Faith PD (and loss of species) in combination with a simultaneous increase in MPD is projected to occur widely across mainland Europe and is especially widespread across Spain (Figure 2), indicating that future species assemblages in these areas are projected to be less related despite the overall decrease in species numbers. Focussing on Spain, one possible explanation for these changes could be that species which currently have their breeding grounds limited to northern Africa are projected to move into Spain under climate change (Barbet-Massin et al., 2010; Guiterrez, 2001), potentially adding species to the assemblages in Spain that are very different from currently occurring species. These interpretations made for the example of Europe can be made for other regions in a similar fashion. In general, our results show that it is important to consider additional biodiversity indices to the usual SR metrics to evaluate potential impacts of the redistribution of species under climate change; in this case, even though the changes in SR are mostly matched by the change in Faith PD, the mismatch with MPD shows that different aspects of biodiversity of a species assemblage can be affected in different ways.

The extinction of species from, and introduction of species into species assemblages can lead to phylogenetic restructuring (Winter et al., 2009). For example, it is well supported that anthropogenic impacts such as habitat alteration or species invasions into assemblages can lead not only to taxonomic but also phylogenetic homogenisation of species communities, through mechanisms of extinction and replacement (McKinney & Lockwood, 1999; Nowakowski et al., 2018; Olden et al., 2018). Interpreting what changes to the phylogenetic structure of species assemblages could mean in conservation terms is difficult. Generally, more phylogenetically diverse communities have been associated with higher ecosystem stability (Cadotte et al., 2012) and the continuity of ecological functions and services (Cadotte et al., 2011). Furthermore, species assemblages with a higher phylogenetic diversity are thought to be more resilient to ecological disturbance (Faith, 1992a). Therefore, those assemblages that maintain a high Faith PD or experience an increase in Faith PD could be less vulnerable and continue to contribute to the delivery of ecosystem services relative to assemblages where PD declines (Faith et al., 2010). A decrease in MPD that leads to increased clustering within species assemblages might also be problematic. As closely related species frequently share similar traits, clustering could increase competition between species within an assemblage (Procheş et al., 2008). Overall, our analysis shows that species range shifts under climate change are not only affecting species numbers but are likely to impact the phylogenetic structure of species assemblages in ways that could affect the stability of communities and also the future provision of ecosystem services and ultimately human wellbeing (Faith et al., 2010; Srivastava et al., 2012).

Looking at the potential impacts of climate change on the phylogenetic structure of species assemblages in more detail we showed that the species assemblages are projected to change into each of four possible directions (Figure 3). We classified these four different directions into four types of potential changes to the phylogenetic structure of species assemblages, which are predominant in different parts of the world (Figure 3). Firstly, we interpreted those assemblages that are undergoing a projected loss in Faith PD with a simultaneous gain in MPD as experiencing increasing phylogenetic over-dispersion, meaning there will be fewer species that are very distantly related (Cavender□Bares et al., 2004; Webb et al., 2002). Phylogenetic over-dispersion can often be observed in ecological communities that are thought to have evolved under competitive exclusion (Cooper et al., 2008; Dehling et al., 2014; Emerson & Gillespie, 2008; Slingsby et al., 2006). For the projections under climate change, this pattern might be largely driven by the loss of species from assemblages, since the increase in phylogenetic over-dispersion occurs widely in species rich, tropical areas that are also projected to undergo high species losses (Fig. 1d and 2b).

Second, we interpreted those assemblages with a projected loss of MPD and gain in Faith PD as experiencing increasing phylogenetic clustering, meaning that they are gaining species but these are mostly related to each other or to the species already occurring in the assemblage. Phylogenetic clustering has been observed in species assemblages that evolved under environmental filtering, which selected for species that are able to cope with the same conditions (Asefa et al., 2017; Emerson & Gillespie, 2008); but see (Cadotte & Tucker, 2017; Kraft et al., 2015). Under our climate change projections this type of compositional change is mainly found at high northern latitudes as well as along the southern Andes and the Tibetan plateau (Fig. 3b). This is in line with our hypothesis that uniform responses across species, like highly directional range shifts (such as widespread shifts towards higher latitudes and altitudes; (Devictor et al., 2008; J Lenoir et al., 2008; Sekercioglu et al., 2008; Virkkala & Lehikoinen, 2014)) might select for species with similar traits and subsequently increase phylogenetic clustering.

Third, we interpreted species assemblages with a projected loss in both MPD and Faith PD as undergoing an overall increase in phylogenetic homogenisation, meaning that they are also experiencing a loss in Faith PD (and hence evolutionary history) whilst the overall relatedness between species slightly decreases, leaving the assemblage with fewer and more closely related species than before. Phylogenetic homogenisation is often observed in species communities that have undergone anthropogenic disturbance, like habitat conversion through urbanisation or agricultural expansion or intensification (Liang et al., 2019; Sol et al., 2017). Under the climate change projections utilized here, phylogenetic homogenisation occurs least frequently (Table 1), mainly across Eastern Europe (Fig. 3b), and is characterized by relatively small changes in all three measures SR, Faith PD and MPD (Fig. 3a). That we do not see drastic losses here could be due to the high mobility of many bird species. Looking at the impacts for other, less mobile, taxa or assuming a no-dispersal scenario would likely increase the number of assemblages in this group due to potentially higher extinction (Foden et al., 2013).

Finally and fourth, we interpreted those assemblages with a projected increase in both MPD and Faith PD as undergoing increasing diversification, meaning that the assemblages are becoming overall more phylogenetically rich with an increase in evolutionary history and a decrease in the overall relatedness of the species. These increases are mostly projected into currently species poor regions such as parts of the North American, European and Asian Taiga and some of the world’s deserts, such as the Saharan, Arabian, Australian and Kalahari deserts (Fig. 3b). This distribution is probably due to the fact that in species-poor regions, the addition of even just one species is highly likely to increase phylogenetic diversity for simple mathematical reasons.

### Projected non-random changes in phylogenetic assemblage structure

The magnitude in which phylogenetic diversity is changing depends on the structure of the phylogenetic tree (Cadotte & Davies, 2016; Heard & Mooers, 2000; Vellend et al., 2011) and, for Faith PD, can mirror or deviate from the taxonomic diversity (Frishkoff et al., 2014). In contrast to earlier studies that projected global extinction risk from climate change to be evenly distributed across the phylogeny for some taxa (Pio et al., 2014; Thuiller et al., 2011), we show that projected changes in the phylogenetic diversity of species assemblages under climate change can differ significantly from equivalent random changes to the species pool (Fig. 4). Looking at the decreases in Faith PD and MPD through the loss of species, we project that for terrestrial bird species there are substantial areas across the globe where phylogenetic diversity decreases or increases more than expected at random.

Especially for Faith PD, the non-random decreases might be connected to the local loss of ancient lineages, since assemblages where Faith PD decreases more than expected at random are mostly located in tropical Africa and South America which host higher numbers of ancient lineages (Procheş et al., 2015; Voskamp et al., 2017). By contrast, assemblages where Faith PD decreases less than expected at random are mostly located at high northern latitudes which are known to host fewer ancient lineages (Procheş et al., 2015). Our findings conflict somewhat with a study that investigated non-random declines in Faith PD under climate change in various mammal, plant and insect families in the Floristic Cape Region of South Africa and found that the observed decline differed little from random simulations (Pio et al., 2014). This difference in the results might reflect variation between taxa, or the fact that the latter analysis was conducted in a biodiversity hotspot that is rich in endemics and old lineages (Cape floristic region has over 1500 plant genera, 30% of which occur nowhere else globally), but is more likely due to the scale of the analysis.

Importantly, aside from investigating potential non-random declines in phylogenetic diversity, we used novel methods to disentangle the impacts of projected assemblage gains and losses under climate change. We found that, in particular, the impacts of assemblage gains differ significantly from random across large areas globally. Our results show that identifying areas where the changes in phylogenetic diversity differ from random is important, since they could be more widespread than previously assumed, and our analyses highlight where projected richness changes might not fully reflect the risk that climate change poses to communities.

When disentangling the impacts of species loss and gain, we identified areas in the world where the impacts of species losses and gains on the phylogenetic diversity of the assemblages are causing changes in the same direction. The outcome could be positive, where the PD or MPD increases, through species entering an assemblage, are significantly higher than expected whilst the PD or MPD decreases, through species being lost from the same assemblage, are significantly lower than expected. Or the outcome could be negative, with lower PD or MPD gains than expected based on species gains and greater PD or MPD declines than expected based on species losses. For such assemblages, the overall changes in phylogenetic diversity will be clearly positive or negative. By contrast, there are also situations where the loss and gain of species have different implications for the assemblage phylogenetic diversity, highlighting the utility of a multi-metric approach to studying impacts. For example, in such areas the species gains can lead to a greater PD or MPD than expected but, at the same time, species losses lead to lower PD or MPD than expected (or vice versa). In such assemblages, the net change in PD or MPD might be close to zero, but the underlying changes in the species composition, are nonetheless high, indicating greater shifts in assemblage composition than would be apparent if gains and losses of species were not separately assessed. Our results show that disentangling the impacts of species gains and losses can aid understanding of how assemblage phylogenetic diversity might be impacted by climate change, and that important aspects of this impact might be masked if only considering overall phylogenetic diversity change or if only focussing on losses.

### Data limitations and model uncertainties

There are several data-related and methodological caveats that need to be considered when interpreting the results of this study. Firstly, we were gridding species distribution range maps at a 0.5° × 0.5° grid (circa 55×55 km) resolution. Aside from the debate around the utility of range maps for SDMs (Herkt et al., 2017), the coarse resolution of the resultant distribution data and the associated climate data used in the modelling may lead to potential misinterpretation of the finer-scale relationships between species occurrences and climate. However, our global assessment for all bird species could only be conducted at this scale due to a paucity of more highly resolved occurrence data, as is common with almost all similar global analyses. Consequently, our study is useful in outlining broad trends in potential assemblage changes but should not be extended to infer locality- or species-specific responses.

Additionally, our projections of species’ current and future ranges are solely based on the climatic niche of a species. Both changes in land cover and land-use as well as biotic interactions will have an impact on the future ranges of the modelled species (Engelhardt et al., 2020; Godsoe & Harmon, 2012; Sirami et al., 2017). Furthermore, biotic interactions will also have an impact on the likelihood of the future establishment of species (Mitchell et al., 2006). Whilst some promising modelling approaches incorporate biotic interactions into SDMs (Kissling et al., 2012), this is still a major challenge (Zurell et al., 2020), especially when working on a global scale and considering entire higher taxa. Similarly, our ability to integrate land-use and climate change remains limited (Sirami et al., 2017). Although there have been advances in the understanding of species-specific habitat suitability (Methorst et al., 2017; Rondinini et al., 2011) and efforts to derive links between land-use change and biodiversity loss (Newbold et al., 2014, 2015), obtaining biologically relevant data at the scale of this study remains a challenge. Nevertheless, despite these sources of uncertainties, we believe our method is robust and highlights broad geographic trends in potential phylogenetic change as well as identifying ways in which both heterogeneous and homogenous responses of species to climate change could impact the phylogenetic structure of species assemblages.

### Implications for conservation

The value and practicability of using phylogenetic diversity indices for conservation purposes has been widely discussed (Winter et al., 2012). Phylogenetic measures can add valuable information for conservation planning (Pollock et al., 2015, 2017), and the preservation of phylogenetic diversity could be key to the resilience of communities to environmental change (Faith, 1992a, 1992b). Among the different actions planned to reduce biodiversity loss and reach the 2050 Vision on Biodiversity, climate change action plays a significant role (Secretariat of the Convention on Biological Diversity, 2020). Although no phylogenetic signal in climate change vulnerability had been identified prior to this study, our results showing strong heterogeneity in assemblage-level changes reinforce the utility of phylogenetic indices for climate impact studies. Whilst our results should not be interpreted at a local scale, they do highlight the implications that climate change induced range shifts could have on the phylogenetic diversity of species assemblages. The widespread projected changes to phylogenetic diversity we demonstrate, indicate a potential for strong changes to the diversity of species’ traits and attributes important for maintaining functioning ecosystems. They thus emphasise the need to minimise future climate change although, for the conservation of biodiversity, climate change mitigation strategies must run alongside reductions in other drivers of biodiversity loss (Leclère et al., 2020).

## Supporting information

Supplementary results and discussion

## Acknowledgements

We thank Dan Rosauer for valuable discussions on the study design and methodology.

## Author contributions

AV and SGW conceived the initial idea. AV, SAF and CH designed the study. AV analyzed the data with input from SAF, CH, MFB and SGW. AV wrote the manuscript with comments from all contributing authors.

## Supplementary material

**Figure S1:** Map of the distribution of the species that had to be excluded from the analysis due to limited range extent or low model performance

**Figure S2:** *Adapted figure 2:* Projected changes in SR Faith PD and MPD for RCP 2.6 under a medium dispersal scenario

**Figure S3:** *Adapted figure 3:* Comparison of phylogenetic assemblage structure for RCP 2.6 under a medium dispersal scenario

**Figure S4:** *Adapted figure 4:* Projected changes in Faith’s phylogenetic diversity (Faith PD) and mean phylogenetic distance (MPD) of species assemblages for RCP 2.6 under a medium dispersal scenario

**Figure S5:** *Adapted figure 2:* Projected changes in SR Faith PD and MPD for RCP 6.0 under a low dispersal scenario

**Figure S6:** *Adapted figure 3:* Comparison of phylogenetic assemblage structure for RCP 6.0 under a low dispersal scenario

**Figure S7:** *Adapted figure 4:* Projected changes in Faith’s phylogenetic diversity (Faith PD) and mean phylogenetic distance (MPD) of species assemblages for RCP 6.0 under a low dispersal scenario

**Table S1:** Species numbers included in the different steps of the analysis

**Table S2:** *Adapted table 1:* The overall terrestrial area, globally and per continent, that falls into the four different change categories for RCP 2.6 under a medium dispersal scenario

**Table S3:** *Adapted table 2:* Combined projected changes in Faith’s phylogenetic diversity and mean phylogenetic distance for RCP 2.6 under a medium dispersal scenario

**Table S4:** *Adapted table 1:* The overall terrestrial area, globally and per continent, that falls into the four different change categories for RCP 6.0 under a low dispersal scenario

**Table S5:** *Adapted table 2:* Combined projected changes in Faith’s phylogenetic diversity and mean phylogenetic distance for RCP 6.0 under a low dispersal scenario

