## Supplementary results and discussion for "Climate change impacts on the phylogenetic diversity of the world’s terrestrial birds: more than species numbers"

### Additional methods

#### Species distribution data

We gridded the range polygons onto a 0.5° x 0.5° latitude–longitude grid, to match the spatial resolution of the climate data. All species whose range polygons overlapped a grid cell were added to that cell's species assemblage. Species that occurred in =< 10 grid cells were excluded from further analysis (896 species), due to the difficulties of modelling species with restricted range size.

To derive pseudo-absences for each species, for use in SDMs, we used a distance-weighted approach to randomly choose absence points beyond a species range edge, in which the likelihood of randomly selecting a point decreases with $\frac{1}{De^2}$ where *De* is the distance from range edge (Hof et al., 2018). Using this method, we drew 10 potential sets of pseudo absences for each species. This approach reduces the risk of selecting points too close to the species range and subsequently truncating the response curve (Barbet-Massin et al., 2012; Thuiller, 2004), as well as the risk of selecting absences too far from the range edge which might contain little relevant information for the model (Anderson & Raza, 2010).

#### Species distribution models

The two model algorithms, we applied, were chosen based on their performance in comparison to other model algorithms (Duque-Lazo et al., 2016; Meynard & Quinn, 2007) as well as to provide a contrast between a non-parametric (GAM) and a machine learning (GBM) method. We identified a set of variables that performed well for a representative subset of the world's terrestrial bird species drawn evenly from across the globe, whilst avoiding variable combinations with >= 0.7 correlation (Hof et al., 2018). The selected variables were temperature seasonality (BIO4), maximum temperature of the warmest month (BIO5), annual precipitation (BIO12) and precipitation seasonality (BIO15).

To reduce spatial autocorrelation in the SDMs we applied a blocking approach, following the methods of Bagchi et al. (2013). We divided the data into sampling units, based on the world's ecoregions. These sampling units were then split into 10 approximately equally sized blocks, with each block representing the full climatic parameter space of the chosen bioclimatic variables. Models were subsequently built on nine blocks and tested on the left-out block. For range restricted species (<50 grid cells) this blocking method does not work, here we applied the commonly used 30:70 split with 10 repeated draws.

We evaluated model performance, based on the model fit, using the area under the curve (AUC; (Fielding & Bell, 1997)). The AUC values were calculated across the 10 fitted models following the cross validation for each of the 10 pseudo- absence sets (100 models per species, per model type). We excluded all species that had an average AUC < 0.7 for the two model types from all further analysis, resulting in a final number of 8269 species (Fig. S1 and Table S1).

| **Analysis step** | **# of species** |
|---|---|
| All terrestrial bird species downloaded (BirdLife taxonomy 2015) | 9538 |
| Restricted range species removed | 896 |
| Species whose ranges were merged whilst matching the taxonomies | 224 |
| Species with low model performance removed | 149 |
| ***Remaining species included in the analysis*** | ***8269*** |

**Table S1:** Numbers of species that were excluded from the analysis due to their restricted range or low model performance.

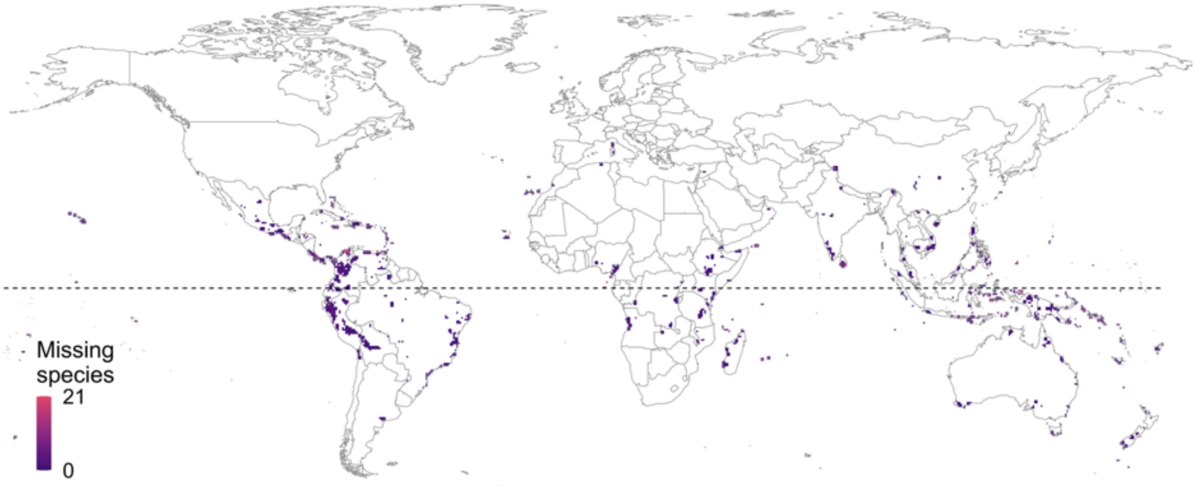


**Fig. S1:** Distribution of the 896 range restricted terrestrial bird species that were excluded from the analysis. Purple indicates low species richness and red indicates high species richness.

#### Additional results

To investigate the robustness of the results, we included an additional lower warming scenario (RCP 2.6) and an additional more restricted dispersal scenario. The results for the lower emission scenario allow to investigate if reducing climate warming would make a difference to the projected changes in the three measures species richness, Faith PD and mean phylogenetic diversity. The extra dispersal scenario allows checking how sensitive the projected changes are to the included dispersal assumption. Including a no dispersal scenario does not make sense for this type of study, because it would not allow for species to be gained by a grid cell and would thus not be comparable. We therefore opted for a restricted dispersal scenario adding a species-specific buffer to the individual species polygons calculated as $\frac{d}{4}$, where d equals the diameter of the largest range polygon of a species (for details see (Hof et al., 2018)).

**Results shown for a low emission pathway (RCP 2.6)**

Here we reproduced the results from the main manuscript (Fig. 2 to 4 and Table 1 and 2) under a low warming scenario (RCP 2.6). Including the additional warming scenario, we found that under a lower emission scenario the projected losses in species richness were overall slightly reduced but the spatial pattern in the projected changes in species richness, Faith PD and MPD were similar Fig. S2. Looking at the four different categories of phylogenetic assemblage structure we found, that the category 'increasing clustering, is projected to be more widespread under the low emission scenario whilst the category 'increasing over-dispersion' is projected to be less widespread globally (with the exception of Australia), compared to the projected changes under a medium warming scenario (RCP 6.0) (Table S2). But the overall spatial distribution of the four categories stayed largely similar (Fig. S3). The spatial pattern of where increases and decreases in Faith PD and MPD are projected to be higher or lower that what we would expect at random stayed remarkably similar (Fig. S4). Importantly, those areas that are projected to undergo significantly higher decreases than random by species disappearing from assemblages, whilst at the same time experiencing significantly lower increases than random through species being gained by assemblages are reduced a lot under the low warming scenario (Table S3).

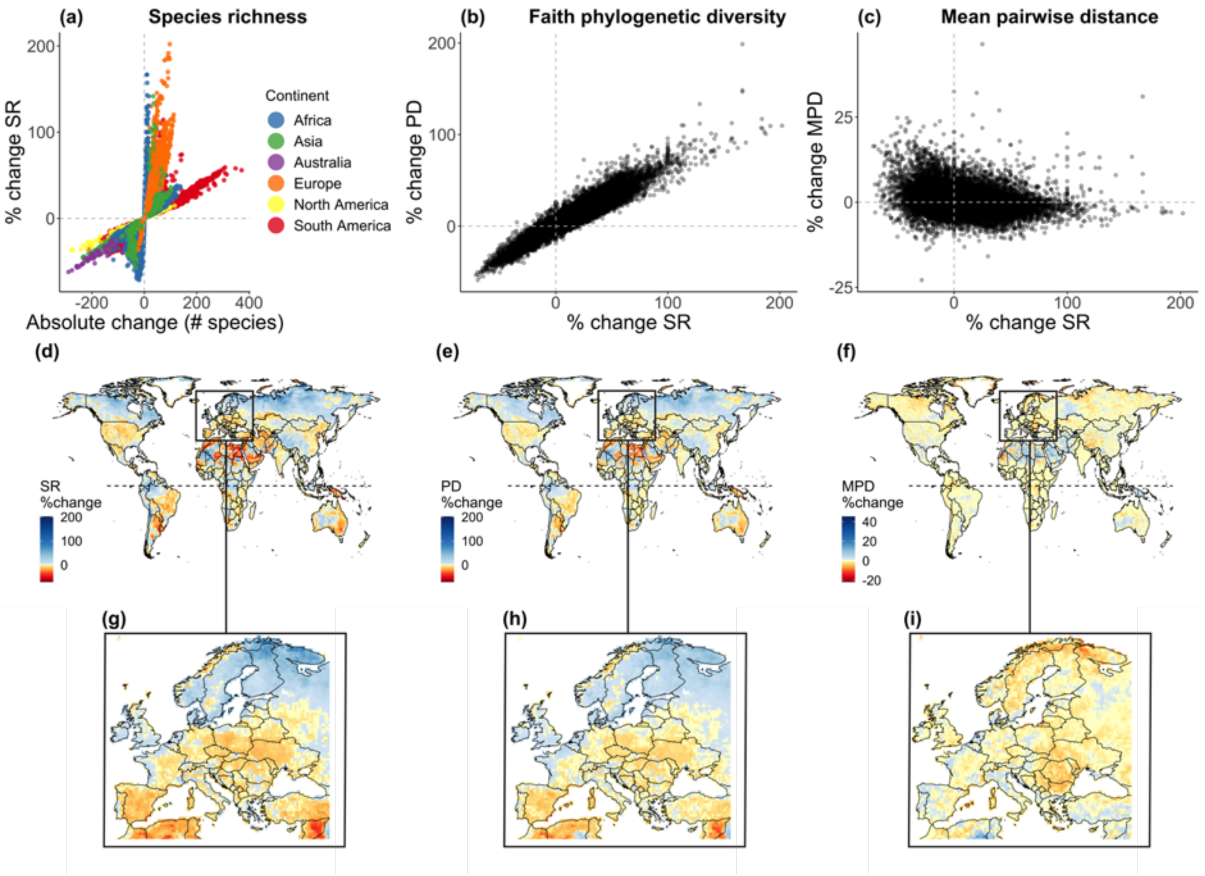


**Fig. S2:** Projected changes in species richness (SR), Faith's phylogenetic diversity (Faith PD) and mean phylogenetic distance (MPD) under a low emission scenario (RCP 2.6) and a medium dispersal scenario by 2080. (a) shows the percentage change in SR against absolute change in SR; (b) the percentage change in Faith PD against percentage change in SR; (c) the percentage change in MPD against percentage change in SR (d) the spatial distribution of percentage change in SR; (e) the spatial distribution of percentage change in Faith PD and (f) the spatial distribution of percentage change in MPD. The percentage change for all three measures is shown in detail for Europe (g – i). Red indicates a negative change (e.g. loss in species richness, Faith PD or MPD), blue indicates a positive change (e.g. gain in species richness, Faith PD or MPD).

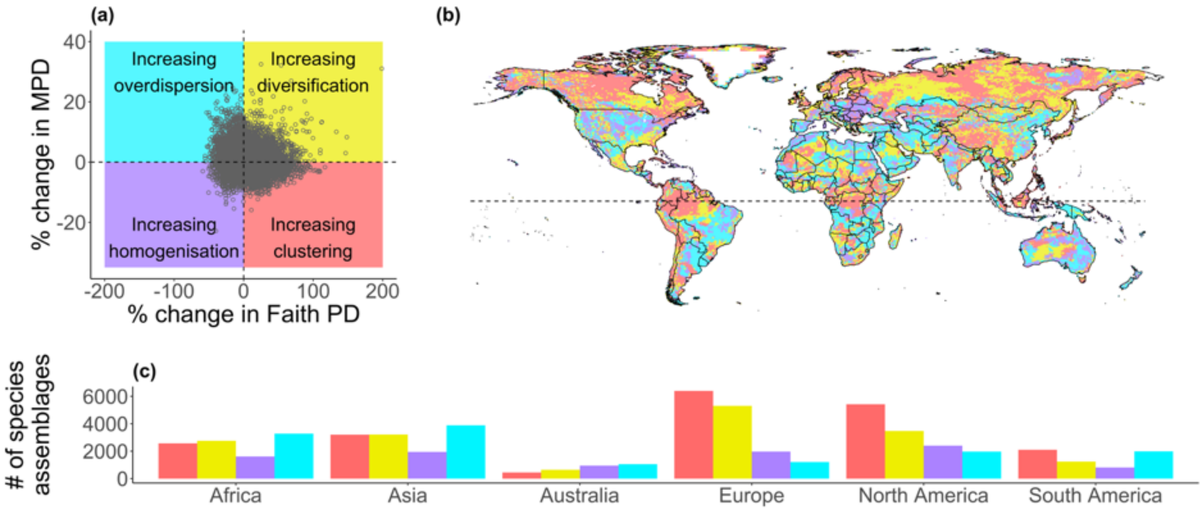


**Fig. S3:** Comparison of phylogenetic assemblage structure as indicated by mean phylogenetic distance (MPD) versus by Faith's phylogenetic diversity (Faith PD) under a low emission scenario (RCP 2.6) assuming a medium dispersal scenario by 2080. The scatterplot (a) shows percentage change in MPD against percentage change in Faith PD, divided into four categories of change using the median along each axis. The map (b) shows the spatial distribution of the species assemblages falling into one of these four categories, and the bar chart (c) shows the number of assemblages per category across different continents. The four defined categories are: grid cells with a projected gain in MPD and loss in Faith PD leading to increasing phylogenetic over-dispersion of these species assemblages (blue); grid cells with a projected loss in both MPD and Faith PD, leading to increasing homogenisation of these species assemblages (purple); grid cells with a projected loss of MPD and gain in Faith PD, indicating increasing phylogenetic clustering of these species assemblages (red); and grid cells with a projected gain in both MPD and Faith PD, indicating increasing diversification within these species assemblages (yellow).

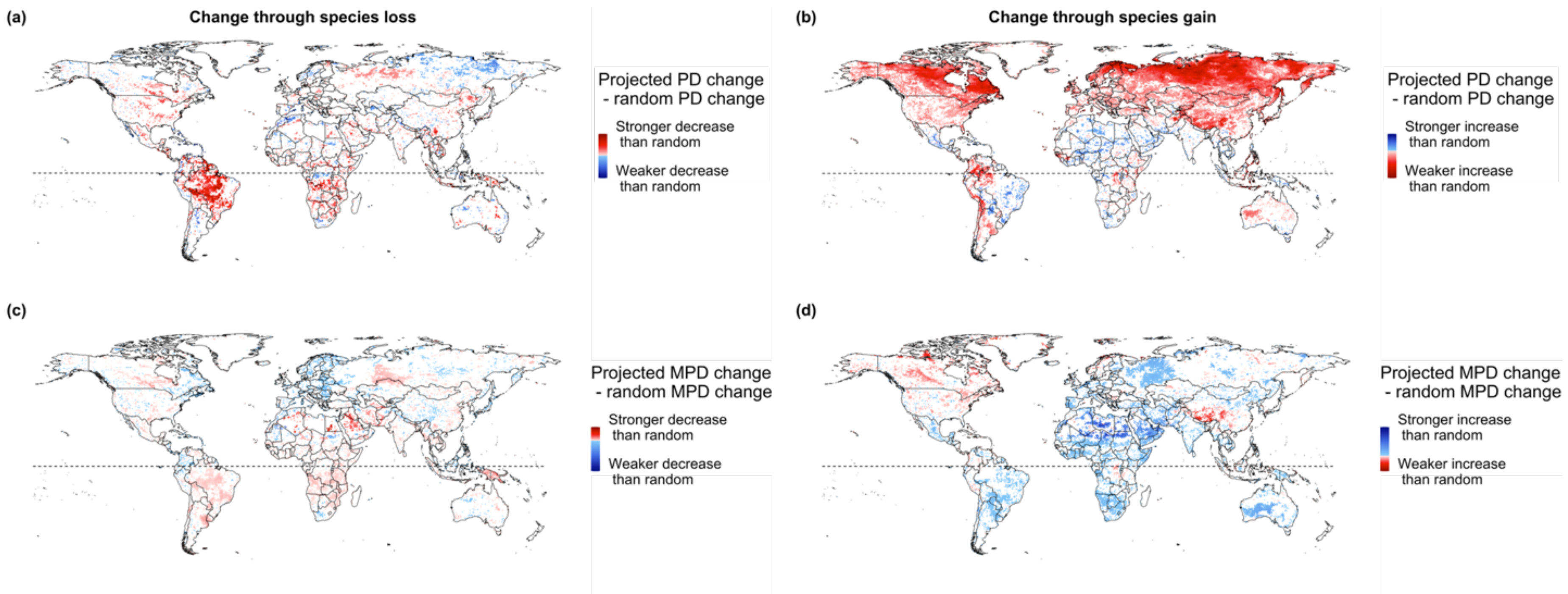


**Fig. S4:** The significance and direction of projected changes in Faith's phylogenetic diversity (Faith PD) and mean phylogenetic distance (MPD) of species assemblages (grid cells), through species that are projected to be lost from (a and c) and gained into (b and d) assemblages, in comparison to expected changes if species were lost and gained at random. Results are shown for a low emission scenario (RCP2.6) and a medium dispersal scenario by 2080. Difference values for species being lost from an assemblage are calculated as shown in Fig 1**.** For the maps of change in Faith PD/MPD through species being lost from an assemblage (a and c), red indicates that the loss of Faith PD/MPD caused by the species that are projected to be lost from the assemblage is significantly higher than what would be expected if the same number of random species would be lost; blue indicates that the loss is significantly lower than what would be expected if random species would be lost (significance is derived using a two-sided p-value < 0.05 or > 0.95). For the maps of change in Faith PD/ MPD through species being gained into an assemblage (b and d), red indicates that the gain in Faith PD/MPD through the species projected to be gained into the assemblage is significantly lower than what would be expected if the same number of random species would be gained into the assemblage, blue indicates that the gain is significantly higher than what would be expected if random species would be gained. A gain or loss in Faith PD signifies a significant increase or decrease in total evolutionary history represented, respectively; a gain or loss in MPD signifies a significant decrease or increase in average relatedness, respectively. White areas in each map have no significant changes compared to random species gain or loss. White areas in each map have no significant changes compared to random species gain or loss.

| | Increasing homogenisation | | Increasing clustering | | Increasing over-dispersion | | Increasing diversification | |
|---|---|---|---|---|---|---|---|---|
| | *Area in km²* | *%* | *Area in km²* | *%* | *Area in km²* | *%* | *Area in km²* | *%* |
| **Africa** | 4,640,503 | 16 | 7,649,200 | 25 | 9,524,786 | 32 | 8,029,771 | 27 |
| **Asia** | 4,940,022 | 16 | 8,147,855 | 26 | 9,849,707 | 32 | 8,105,373 | 26 |
| **Australia** | 2,608,488 | 31 | 1,226,697 | 14 | 2,894,298 | 34 | 1,747,884 | 21 |
| **Europe** | 3,373,158 | 13 | 9,069,611 | 43 | 2,203,352 | 8 | 8,066,093 | 36 |
| **North America** | 4,445,394 | 18 | 8,081,974 | 41 | 4,095,858 | 15 | 6,160,564 | 26 |
| **South America** | 2,289,488 | 13 | 6,102,738 | 34 | 5,627,231 | 32 | 3,580,304 | 20 |
| **Global km²** | 22,297,053 | | 40,278,075 | | 34,195,232 | | 35,689,989 | |
| **Global %** | 16 | | 34 | | 22 | | 28 | |

**Table S2:** The overall terrestrial area, globally and per continent, that falls into the four different categories of combined change in two phylogenetic structure metrics, Faith's phylogenetic diversity (Faith PD) and mean phylogenetic distance (MPD) (as shown in Fig. 3): increasing homogenisation (loss of PD and MPD); Increasing clustering (gain in PD and loss of MPD); Increasing over-dispersion (loss of PD and gain in MPD) and Increasing diversification (gain in PD and MPD). The extent of the area projected to fall into the four different categories is derived assuming a low emission scenario (RCP 2.6) and a medium dispersal scenario by 2080. The area extent is given in $km^2$ as well as in the percentage of the total terrestrial area, per continent and globally.

| Combinations of significantly non-random changes | | Faith PD | | MPD | |
|---|---|---|---|---|---|
| | *Continent* | *Area in km²* | *%* | *Area in km²* | *%* |
| **More decrease than under random species loss AND Less increase than under random species gain** | ***Africa*** | 489,741 | 0.16 | 235,602 | 0.75 |
| | ***Asia*** | 951,169 | 3.27 | 136,322 | 0.42 |
| | ***Australia*** | 62590 | 0.72 | 5,160 | 0.07 |
| | ***Europe*** | 2,022,452 | 8.83 | 21,812 | 0.12 |
| | ***North America*** | 1,616,817 | 7.46 | 444172 | 2.25 |
| | ***South America*** | 832,329 | 4.50 | 59526 | 0.33 |
| | ***Global*** | 5,975,096 | 5.29 | 902,595 | 0.78 |
| **More decrease than under random species loss AND More increase than under random species gain** | ***Africa*** | 3,854,304 | **12.76** | 5,472,936 | **17.93** |
| | ***Asia*** | 2,823,905 | 8.96 | 3,987,098 | **12.75** |
| | ***Australia*** | 395,301 | 4.58 | 622349 | 7.09 |
| | ***Europe*** | 2,312,917 | 9.91 | 1,750,800 | 7.44 |
| | ***North America*** | 2,972,783 | **14.14** | 2,538,927 | **12.91** |
| | ***South America*** | 5,120,569 | **27.87** | 3,834,597 | **21.54** |
| | ***Global*** | 17,479,780 | **12.71** | 18,206,707 | **12.96** |
| **Less decrease than under random species loss AND More increase than random species gain** | ***Africa*** | 974,407 | 3.36 | 615,355 | 2.15 |
| | ***Asia*** | 1,004,037 | 3.18 | 1,724,916 | 5.54 |
| | ***Australia*** | 282,466 | 3.28 | 220,505 | 2.57 |
| | ***Europe*** | 1,620,078 | 8.38 | 2,111,098 | 8.79 |
| | ***North America*** | 640,296 | 3.00 | 1,361,483 | 5.48 |
| | ***South America*** | 455,989 | 2.68 | 577,476 | 3.18 |
| | ***Global*** | 4,977,273 | 4.12 | 6,610,834 | 5.36 |
| **Less decrease than under random species loss AND Less increase than under random species gain** | ***Africa*** | 94,893 | 0.32 | 3076 | 0.01 |
| | ***Asia*** | 400,490 | 1.32 | 400589 | 1.24 |
| | ***Australia*** | 19,089 | 0.23 | 5479 | 0.07 |
| | ***Europe*** | 1,381,674 | **7.22** | 94,877 | 0.38 |
| | ***North America*** | 364,947 | 1.72 | 303,773 | 1.15 |
| | ***South America*** | 130,579 | 0.75 | 58135 | 0.31 |
| | ***Global*** | 2,391,671 | 2.59 | 865,927 | 0.64 |

**Table S3:** Combined projected changes in Faith's phylogenetic diversity (Faith PD) and mean phylogenetic distance (MPD), indicating the proportions of those assemblages across the globe where each measure changes significantly compared to both, a randomized gain and a randomized loss of species (as shown in Fig. 4). The extent of the area projected to fall into the four different combinations is derived assuming a low emission scenario (RCP 2.6) and a medium dispersal scenario by 2080. The area extent is given in km$^2$ as well as in terms of percentage of the total global land mass; percentage values above 10% are printed in bold.

**Results shown for a medium emission pathway (RCP 6.0) and a low dispersal scenario**

Here we reproduced the results from the main manuscript (Fig. 2 to 4 and Table 1 and 2) assuming a low dispersal scenario (calculating the dispersal buffer as $\frac{d}{4}$, where d equals the diameter of the largest range polygon of a species). Including the additional dispersal scenario, showed that the overall results are robust to changing the dispersal assumption, despite the effect that dispersal has on projected species richness patterns (Hof et al., 2018). The losses in species richness and decreases in Faith PD and mean pairwise distance (MPD) were, as expected, higher under a restricted dispersal scenario but the observed patterns remained stable (Fig. S5). The distribution of the four categories of change in the phylogenetic assemblage structures remained overall largely similar, with increases in the category 'increased clustering' and decreases in the class 'increasing over-dispersion' (Fig. S6 and Table S4). The spatial pattern of where decreases and increases in Faith PD and MPD are projected to be higher or lower than what we would expect at random were largely similar under a low dispersal scenario. This is with the exception of areas where assemblages are projected to increase less in MPD than expected through the gain of species, these were reduced across the Palearctic and Nearctic (Fig. S7 and Table S5). This is probably due to less species being projected to shift as far northwards under a more restricted dispersal assumption.

Overall, we find that changes in the estimated dispersal ability will affect the strength of the projected changes but not the projected spatial pattern and directions of change.

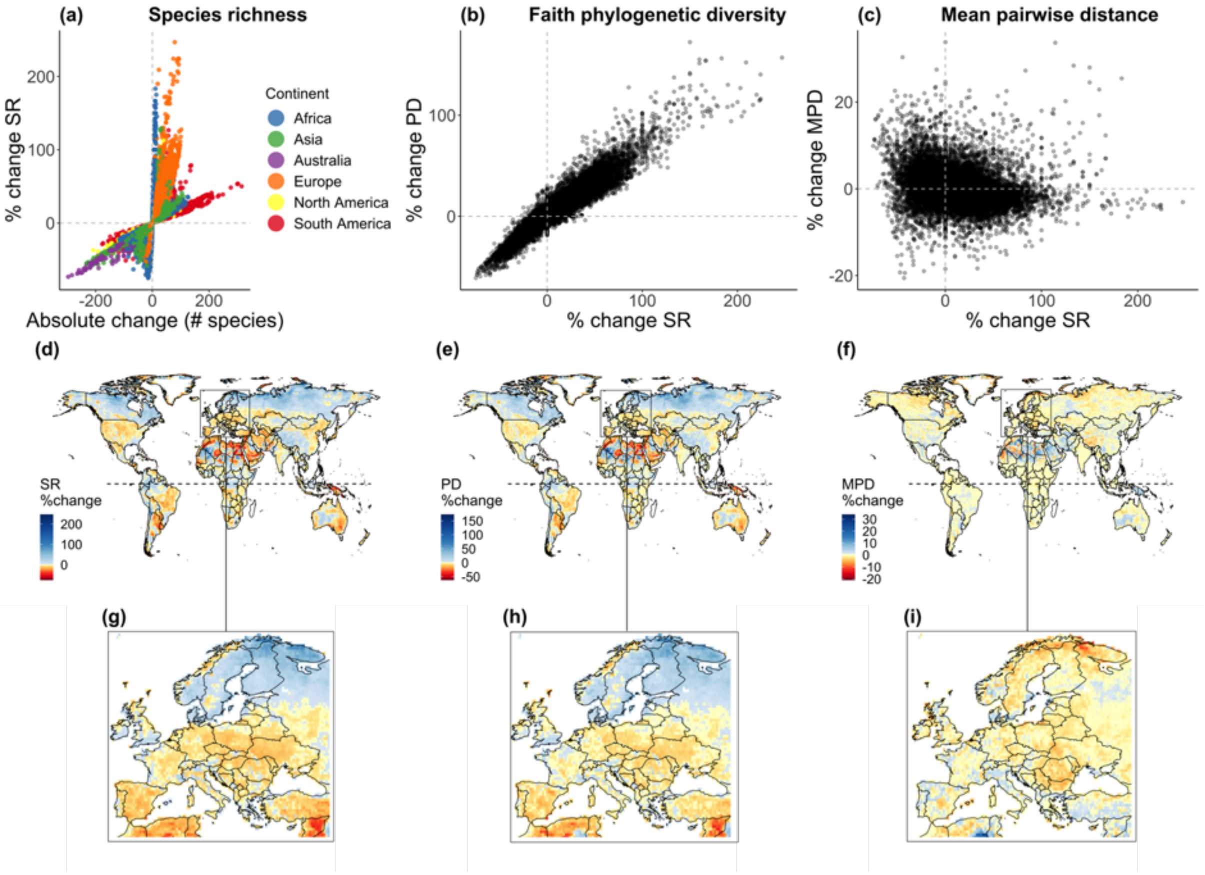


**Fig. S5:** Projected changes in species richness (SR), Faith's phylogenetic diversity (Faith PD) and mean phylogenetic distance (MPD) under a medium emission scenario (RCP6.0) and a low dispersal scenario by 2080. (a) shows the percentage change in SR against absolute change in SR; (b) the percentage change in Faith PD against percentage change in SR; (c) the percentage change in MPD against percentage change in SR (d) the spatial distribution of percentage change in SR; (e) the spatial distribution of percentage change in Faith PD and (f) the spatial distribution of percentage change in MPD. The percentage change for all three measures is shown in detail for Europe (g – i). Red indicates a negative change (e.g. loss in species richness, Faith PD or MPD), blue indicates a positive change (e.g. gain in species richness, Faith PD or MPD).

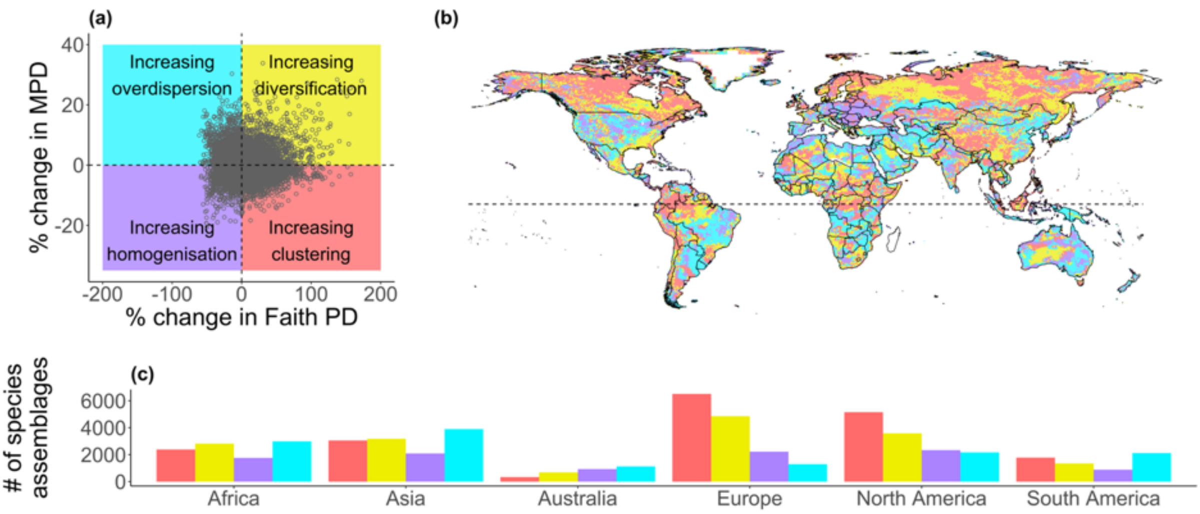


**Fig. S6:** Comparison of phylogenetic assemblage structure as indicated by mean phylogenetic distance (MPD) versus by Faith's phylogenetic diversity (Faith PD) under a medium emission scenario (RCP 6.0) assuming a low dispersal scenario by 2080. The scatterplot (a) shows percentage change in MPD against percentage change in Faith PD, divided into four categories of change using the median along each axis. The map (b) shows the spatial distribution of the species assemblages falling into one of these four categories, and the bar chart (c) shows the number of assemblages per category across different continents. The four defined categories are: grid cells with a projected gain in MPD and loss in Faith PD leading to increasing phylogenetic over-dispersion of these species assemblages (blue); grid cells with a projected loss in both MPD and Faith PD, leading to increasing homogenisation of these species assemblages (purple); grid cells with a projected loss of MPD and gain in Faith PD, indicating increasing phylogenetic clustering of these species assemblages (red); and grid cells with a projected gain in both MPD and Faith PD, indicating increasing diversification within these species assemblages (yellow).

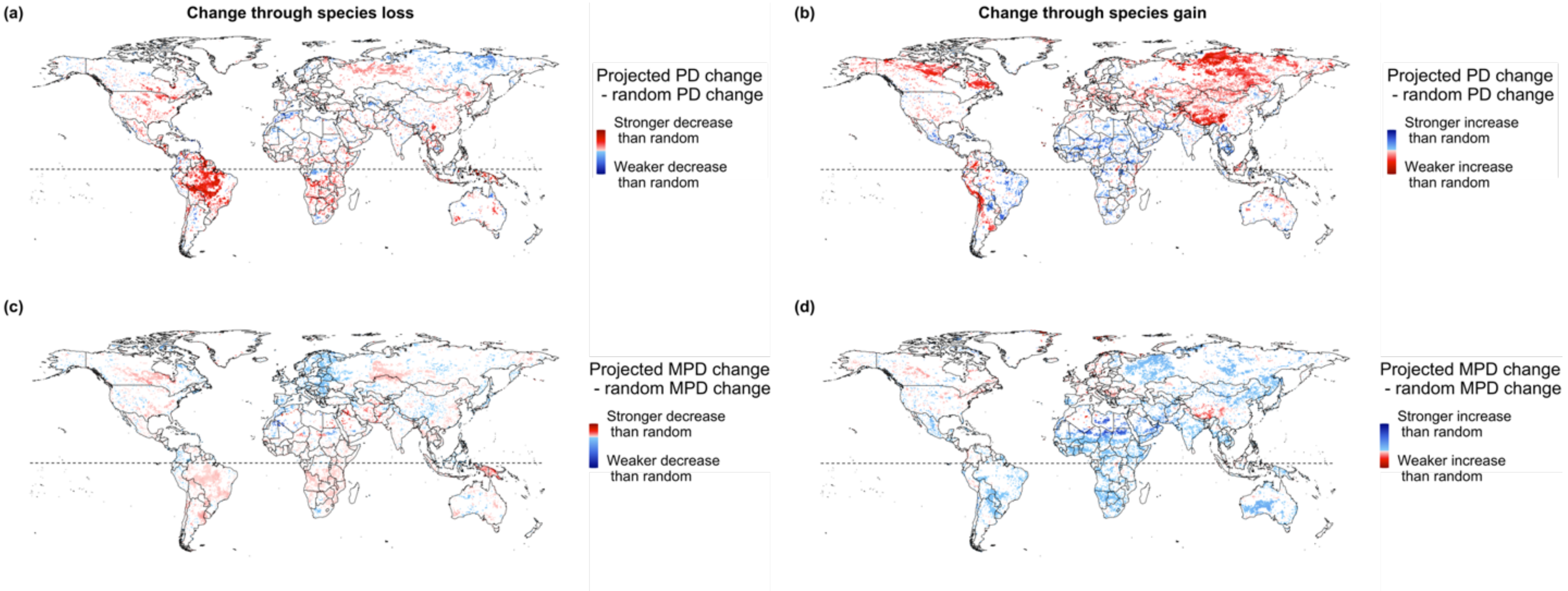


**Fig S7:** The significance and direction of projected changes in Faith's phylogenetic diversity (Faith PD) and mean phylogenetic distance (MPD) of species assemblages (grid cells), through species that are projected to be lost from (a and c) and gained into (b and d) assemblages, in comparison to expected changes if species were lost and gained at random. Results are shown for a medium emission scenario (RCP6.0) and a low dispersal scenario by 2080. Difference values for species being lost from an assemblage are calculated as shown in Fig 1**.** For the maps of change in Faith PD/MPD through species being lost from an assemblage (a and c), red indicates that the loss of Faith PD/MPD caused by the species that are projected to be lost from the assemblage is significantly higher than what would be expected if the same number of random species would be lost; blue indicates that the loss is significantly lower than what would be expected if random species would be lost (significance is derived using a two-sided p-value $< 0.05$ or $> 0.95$). For the maps of change in Faith PD/ MPD through species being gained into an assemblage (b and d), red indicates that the gain in Faith PD/MPD through the species projected to be gained into the assemblage is significantly lower than what would be expected if the same number of random species would be gained into the assemblage, blue indicates that the gain is significantly higher than what would be expected if random species would be gained. A gain or loss in Faith PD signifies a significant increase or decrease in total evolutionary history represented, respectively; a gain or loss in MPD signifies a significant decrease or increase in average relatedness, respectively. White areas in each map have no significant changes compared to random species gain or loss. White areas in each map have no significant changes compared to random species gain or loss.

| | Increasing homogenisation | | Increasing clustering | | Increasing over-dispersion | | Increasing diversification | |
|---|---|---|---|---|---|---|---|---|
| | *Area in km²* | *%* | *Area in km²* | *%* | *Area in km²* | *%* | *Area in km²* | *%* |
| **Africa** | 5,021,187 | 18 | 7,079,835 | 24 | 8,687,174 | 30 | 8,237,073 | 28 |
| **Asia** | 5,303,659 | 17 | 7,754,953 | 25 | 9,860,479 | 32 | 8,036,751 | 26 |
| **Australia** | 2,549,428 | 30 | 919,147 | 11 | 3,096,127 | 37 | 1,880,576 | 22 |
| **Europe** | 3,930,259 | 15 | 9,085,077 | 44 | 2,314,270 | 9 | 7,376,128 | 33 |
| **North America** | 4,117,212 | 18 | 7,824,762 | 39 | 4,284,301 | 16 | 6,484,740 | 27 |
| **South America** | 2,498,837 | 14 | 5,168,894 | 29 | 5,995,953 | 35 | 3,936,077 | 22 |
| **Global km²** | 23,420,582 | | 37,832,667 | | 34,238,304 | | 35,951,345 | |
| **Global %** | 17 | | 32 | | 23 | | 28 | |

**Table S4:** The overall terrestrial area, globally and per continent, that falls into the four different categories of combined change in two phylogenetic structure metrics, Faith's phylogenetic diversity (Faith PD) and mean phylogenetic distance (MPD) (as shown in Fig. 3): increasing homogenisation (loss of PD and MPD); Increasing clustering (gain in PD and loss of MPD); Increasing over-dispersion (loss of PD and gain in MPD) and Increasing diversification (gain in PD and MPD). The extent of the area projected to fall into the four different categories is derived assuming a medium emission scenario (RCP6.0) and a low dispersal scenario by 2080. The area extent is given in $km^2$ as well as in the percentage of the total terrestrial area, per continent and globally.

| Combinations of significantly non-random changes | | Faith PD | | MPD | |
|---|---|---|---|---|---|
| | *Continent* | *Area in km²* | *%* | *Area in km²* | *%* |
| More decrease than under random species loss AND Less increase than under random species gain | ***Africa*** | 166744 | 0.56 | 544482 | 1.82 |
| | ***Asia*** | 592733 | 1.98 | 752435 | 2.31 |
| | ***Australia*** | 56929 | 0.65 | 201319 | 2.42 |
| | ***Europe*** | 754862 | 3.30 | 442346 | 2.00 |
| | ***North America*** | 342943 | 1.99 | 1560515 | 7.13 |
| | ***South America*** | 392674 | 2.19 | 653215 | 3.70 |
| | ***Global*** | 2306885 | 2.03 | 4154312 | 3.37 |
| More decrease than under random species loss AND More increase than under random species gain | ***Africa*** | 3586507 | **12.27** | 4820565 | **16.33** |
| | ***Asia*** | 2737923 | 8.64 | 5142440 | **16.66** |
| | ***Australia*** | 548046 | 6.43 | 1367983 | **16.09** |
| | ***Europe*** | 2751993 | **12.35** | 2749898 | **12.40** |
| | ***North America*** | 3560127 | **17.83** | 5788342 | **26.65** |
| | ***South America*** | 4896792 | **26.78** | 4974197 | **28.07** |
| | ***Global*** | 18081389 | **13.98** | 24843424 | **18.91** |
| Less decrease than under random species loss AND More increase than random species gain | ***Africa*** | 1051508 | 3.75 | 1712927 | 6.13 |
| | ***Asia*** | 913693 | 2.93 | 3778052 | **12.28** |
| | ***Australia*** | 265159 | 3.10 | 791164 | 9.30 |
| | ***Europe*** | 1391729 | 7.13 | 4585997 | **18.72** |
| | ***North America*** | 486853 | 2.14 | 2960371 | **12.26** |
| | ***South America*** | 443754 | 2.56 | 1297934 | 7.33 |
| | ***Global*** | 4552698 | 3.91 | 15126446 | **12.19** |
| Less decrease than under random species loss AND Less increase than under random species gain | ***Africa*** | 55713 | 0.21 | 175269 | 0.62 |
| | ***Asia*** | 252612 | 0.83 | 1400445 | 4.44 |
| | ***Australia*** | 2996 | 0.03 | 215636 | 2.51 |
| | ***Europe*** | 577453 | 2.96 | 1243694 | 4.69 |
| | ***North America*** | 120312 | 0.55 | 886096 | 3.42 |
| | ***South America*** | 102436 | 0.57 | 328483 | 1.81 |
| | ***Global*** | 1111523 | 1.13 | 4249623 | 3.27 |

**Table S5:** Combined projected changes in Faith's phylogenetic diversity (Faith PD) and mean phylogenetic distance (MPD), indicating the proportions of those assemblages across the globe where each measure changes significantly compared to both, a randomized gain and a randomized loss of species (as shown in Fig. 4). The extent of the area projected to fall into the four different combinations is derived assuming a medium emission scenario (RCP6.0) and a low dispersal scenario by 2080. The area extent is given in km$^2$ as well as in terms of percentage of the total global land mass; percentage values above 10% are printed in bold.

### Additional discussion

##### Spatial patterns in the projected change of the phylogenetic measures and non-random changes

The observed opposite patterns in SR/PD and MPD described in Figure 2 are corroborated by the projected non-random changes in PD and MPD. Vast areas in the northern latitudes (e.g. in Europe Scandinavia and parts of the UK, Fig 2h-i) are projected to experience large decreases in Faith PD but moderate to strong increases in MPD, and those areas also contain high percentages of assemblages where those changes are significantly different from those expected if species loss and gain were random (Fig. 4). In particular, most of the northern latitudes are projected to experience major species reshuffling, indicated by significantly stronger decrease in Faith PD than under random species loss (particularly in western North America, Fig. 4a) and simultaneous significantly stronger increase in Faith PD than under random species gain. MPD results corroborate this, as assemblages in northern latitudes experience strong and significant changes in average relatedness through both species loss and gain, but the patterns are often different in Eurasia and North America.

##### Caveats related to the phylogenetic data

Due to computational limitations, we worked with a consensus phylogenetic tree which introduced some uncertainty, compared to using a high number of individual trees where it would be possible to quantify the sensitivity of the projected changes to changes in the underlying phylogenetic tree. Nevertheless, our chosen number of 150 trees is well above the recommended limit to derive a consensus tree from this particular phylogeny (Jetz et al., 2012).
